# SOX2 empowers a rapid tumorigenic programme from the tumour-resistant population in the skin

**DOI:** 10.1101/2025.01.23.634502

**Authors:** Patricia P. Centeno, Christopher Chester, Catriona A. Ford, Patrizia Cammareri, Gareth J. Inman, Thomas Jamieson, Rachel A Ridgway, Richard Marais, Andrew D. Campbell, Owen J. Sansom

**Affiliations:** Cancer Research UK Scotland Institute, Glasgow, UK; Cancer Research UK Manchester Institute, Manchester, UK; School of Cancer Sciences, University of Glasgow, UK

## Abstract

The skin undergoes continuous renewal, relying on stem and progenitor cells to maintain its barrier function. Over time, multiple oncogenic and tumour suppressor events accumulate. How the skin tolerates these mutations in different populations and how cutaneous squamous cell carcinoma (cSCC) emerges from mutant clones is poorly understood. Rapid cSCC development can ensue in melanoma patients treated with BRAF inhibitors (BRAFi) through paradoxical activation of RAF signalling. To model this in mice, we induced MAPK hyperactivation in two epidermal basal populations: stem-cell-like (K14/K5+) and differentiation-committed (IVL+) cells. Remarkably rapid tumorigenesis ensued from the K14/K5+ basal population defining them as a tumour primed. In contrast, despite mutations being retained and populating the skin, IVL+ cells exhibited tumour resistance with long latency. Nevertheless, both populations can act as cSCC cells of origin. Once transformed, they share histological and transcriptomic hallmarks of epidermal transformation independently of the cell-of-origin or the oncogenic driving mutations. Critically, the pioneer factor SOX2 was uniquely upregulated in tumours arising from the resistant population and was both necessary and sufficient for their transformation. Notably, SOX2 bestow an embryonic profile to human keratinocytes and is expressed in ∼20% of human cSCC, suggesting that it could mark tumours originating from committed progenitors.

**Highlights:**

- **Paradoxical MAPK signalling activation is a tumour promoter in cSCC**
- **K14/K5+ represent a tumour-primed population and IVL+ represents a tumour-resistant population, but both can initiate skin cancer**
- **Oncogene-expressing tumour-resistant population takes over the entire skin epidermis but preserves homeostasis**
- **SOX2 is sufficient to initiate cSCC from the tumour-resistant population by preventing delamination and differentiation and promoting stemness**

---

Adult stem cells and progenitors continuously regenerate and maintain the mammalian epidermis, ensuring its structural integrity and protection against the external environment. However, the ageing skin accumulates a high mutational burden, including mutations in known drivers of cutaneous squamous cell carcinoma (cSCC) as well as passenger mutations ^1^. The proliferation of these mutated cells is confined under normal conditions by the skin hierarchy and spatial compartmentalisation, but stressors and promoter events such as wounding, inflammation, UV exposure, and certain oncogenic mutations can overcome these constraints leading to tumorigenesis ^2–7^. cSCC originates from the keratinocytes, and data from murine model systems have illustrated the tumour’s ability to form from different origins, including the hair follicle, the sebaceous glands, and the interfollicular epidermis compartments ^2,8–10^.

The most widely used cSCC mouse model is the well-established two-stage topical application of the chemical carcinogen 7,12-dimethylbenz[a]anthracene (DMBA), followed by the promoting agent 12-O-tetradecanoylphorbol-13-acetate (TPA) ^2,11^. These tumours develop rapidly from the *Lgr6+* stem cells residing in the hair follicle and are predominantly driven by *Hras*^Q61L^ and *Trp53* mutations ^2,12^. Nevertheless, mutations in *HRAS* are relatively uncommon in spontaneous human cSCC (9-16%) but exist in healthy skin as passenger mutations ^1,10,13^. However, HRAS-driven cSCC is commonly found in melanoma patients arising within weeks of treatment with BRAF^V600E^ inhibitors (BRAFi) ^13–15^. This has been attributed to the paradoxical activation of the MAPK signalling pathway by BRAFi, due to the *HRAS* mutations already present in the skin ^15,16^. Spontaneous human cSCC exhibits high mutational burden, mutational heterogeneity and copy number alterations (Bailey et al., 2023), with the main driver mutations concentrated in the Notch (79.5%) and TP53 (71%) signalling pathways ^3^. However, single mutations in these pathways are insufficient to induce cSCC in mice, and additional combinations of genetic driver alterations involving the TGFβ and MAPK signalling pathways are required for the transformation and modelling of disease progression ^8,10,17,18^. The complexity and slow tumour progression of these models have hindered efforts in the field to understand the biology of skin carcinogenesis.

Two models have been proposed in the interfollicular epidermis to explain the basal layer heterogeneity. The first model described the existence of two distinct basal progenitors that contribute differently during homeostasis, wound healing ^19^ and mechanical stretching ^6^. Upon wounding, these progenitors sense reduced cell density and enhance proliferation by undergoing symmetric self-renewing division, whilst the skin adopts a transient fluid-like state to facilitate progenitor migration for regeneration and remodelling ^20^. An alternative second model proposes that the heterogeneity of the basal layer is caused by different levels of commitment to differentiation, suggesting that basal cells continuously transition towards differentiation following cues from the surrounding environment^21^. Nevertheless, the comparison between these two different basal populations in response to oncogenic stimuli remains unexplored.

Here, we assessed the mechanisms that confer competence to initiate cSCC from the two basal cell populations. We generated a suite of mouse models expressing different oncogenes targeted to the basal populations of the interfollicular epidermis, marked by the expression of KRT14+/KRT5+ and IVL+, that rapidly model and resemble human cSCC. Particularly, we modelled the aetiology of fast-growing cSCC in melanoma patients driven by the hyperactivation of the MAPK signalling pathway. We defined a tumour-resistant and a tumour-primed population and uncovered a shared transcriptional programme responsible for transformation. Importantly, we identified SOX2 as a critical factor for the transformation of the tumour-resistant population. SOX2, highly expressed in ∼25% of human cSCC, is known to induce stem cell characteristics as a super pioneer factor ^22^ and is recognised for its ability to reprogram terminally differentiated somatic cells into induced pluripotent stem cells. Our findings demonstrate the different susceptibility of basal populations to transformation revealing that tumours retain the transcriptional priming of their cell of origin.

## Results

### Paradoxical MAPK signalling activation is a tumour promoter in cSCC

To study the susceptibility of the two different basal populations to transformation and tumour formation, we set to generate novel genetically engineered moused models of cSCC that resembled human disease. Patients of cSCC often carry mutations in the *HRAS* gene causing hyperactivation of MAPK signalling. Thus, we expressed *Hras*^G12V^ allele under its endogenous promoter in the basal stem-like population and the basal committed population by using tamoxifen-inducible Cre recombinase expressed from the *Krt14* promoter (*Krt14*-CreERT2; hereafter K14) and the *Ivl* promoter (Ivl-CreERT; hereafter Ivl) respectively (**Figure 1A**).

**Figure 1.**
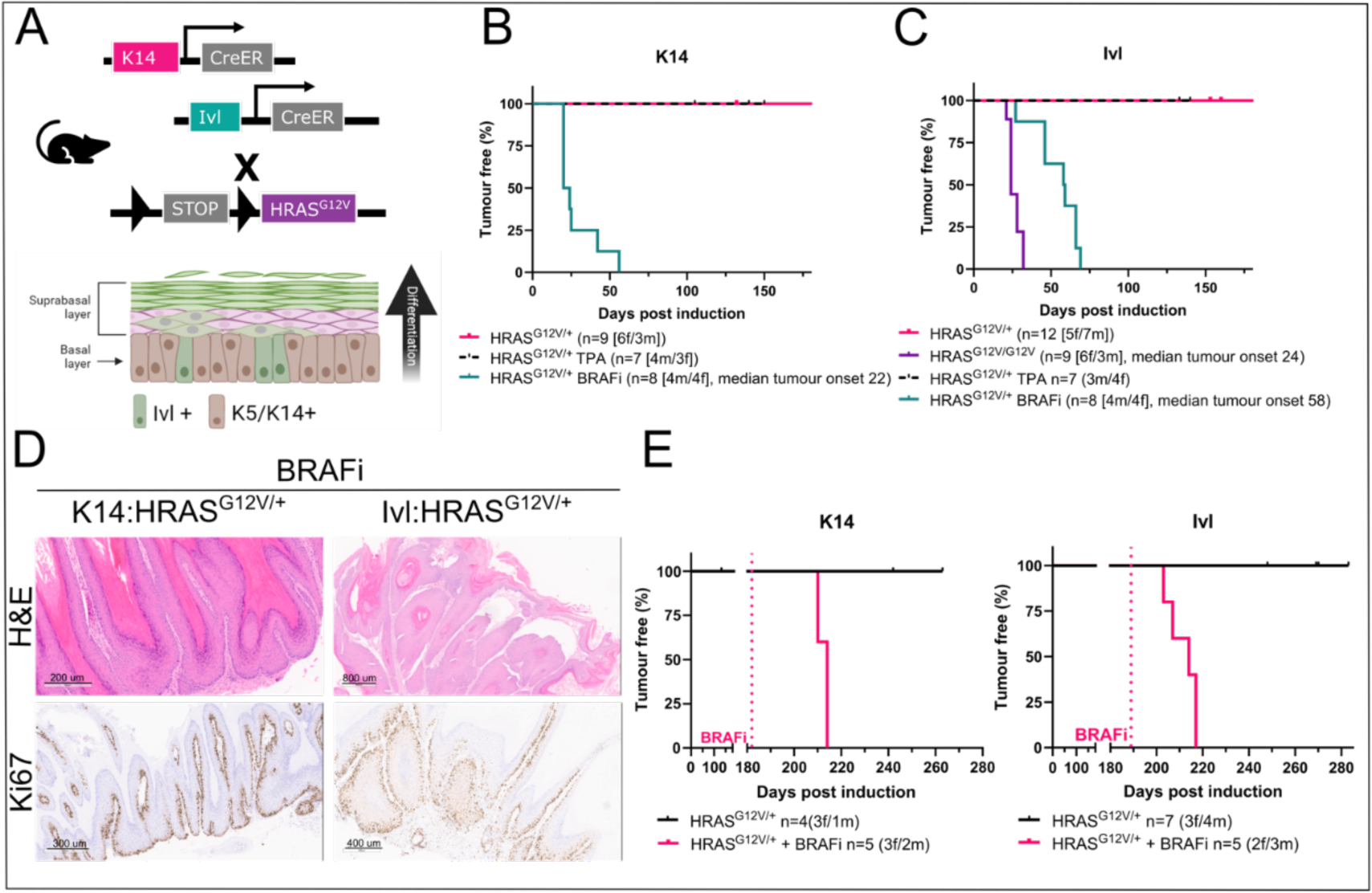
Paradoxical MAPK signalling activation is a tumour promoter in cSCC. (A) Schematic representation of the mouse models used (top) and the epidermis (bottom) formed by the suprabasal, formed by IVL+ cells, and the basal layer, which includes a mixture of K14+/K5+ and IVL+ cells. Created with BioRender. (B-C) Kaplan-Meier tumour-free survival plot for K14:HRAS^G12V/+^ and those treated with the BRAFi Dabrafenib and TPA (B), and Ivl:HRAS^G12V/+^, Ivl:HRAS^G12V/G12V^ and those treated with BRAFi Dabrafenib and TPA (C). (D) Representative histological images including H&E and Ki67 immunohistochemistry (IHC) in tumours derived from K14:HRAS^G12V/+^ and Ivl:HRAS^G12V/+^ mice treated with the BRAFi Dabrafenib at clinical endpoint. (E) Kaplan-Meier tumour-free survival plots for K14:HRAS^G12V/+^ treated with BRAFi Dabrafenib 182 days after oncogene induction (left) and Ivl:HRAS^G12V/+^ treated with BRAFi Dabrafenib 189 days after oncogene induction (right).

Expression of one copy of the *Hras*^G12V^ allele (hereafter HRAS^G12V^) in the K14+ or the IVL+ population did not result in tumorigenesis (**Figures 1B and 1C**). This result is consistent with human studies demonstrating that normal-looking skin carries a battery of mutations, including oncogenes, and that a promotion event is required to trigger transformation^1,2^. Therefore, we topically treated the models with the classical inflammation and tumour-promoting agent known to stimulate epidermal proliferation, TPA, (three times per week) but was insufficient for transformation.

Interestingly, melanoma patients commonly treated with BRAFi, such as Dabrafenib, have an increased risk of developing cSCC due to the presence of dormant *HRAS* mutations in the skin (Hauschild et al., 2012; Oberholzer et al., 2012; Su et al., 2012). Consequently, these patients develop fast-growing tumours within weeks of treatment, driven by the paradoxical activation of the MAPK signalling pathway. Thus, we treated Ivl:HRAS^G12V/+^ and K14:HRAS^G12V/+^ mice with the BRAFi Dabrafenib three times a week to mimic the patient dosing regimen and promote tumorigenesis. Indeed, this resulted in tumorigenesis with a tumour-free survival rate of 22 days for the K14:HRAS^G12V/+^ model and 58 days for the Ivl:HRAS^G12V/+^ model (**Figures 1B and 1C**). BRAFi-promoted tumours exhibited the classic cSCC hallmarks, including keratin pearls, parakeratosis, and nuclear dysplasia (**Figure 1D**). Moreover, we also treated with BRAFi healthy-looking mice carrying one copy of the HRAS^G12V^ in the K14+ and IVL+ mutations at 180±5 days post-oncogene induction. These aged-treated mice rapidly developed tumours within days from the BRAFi treatment, which acts as a potent tumour-promoting agent and mimics the onset of cSCC from melanoma patients carrying *HRAS* mutations (**Figure 1E**).

Two copies of the *Hras*^G12V^ were sufficient to drive tumorigenesis without the need for a promoting agent due to higher levels of MAPK activation. Specifically, two copies of *Hras*^G12V^ expressed in the K14+ population resulted in mice becoming unwell, probably due to the critical function of HRAS in the gastrointestinal epithelium before any skin-related malignancies appeared. Nevertheless, K14:HRAS^G12V/G12V^ mice showed hyperproliferation of the skin basal layer, evidenced by increased Ki67 expression (**Figure S1**). In the IVL+ basal population, two copies of the allele drove tumorigenesis with a tumour-free survival of 24 days (**Figures 1C and S1**). The resulting tumours displayed similar histopathological features (**Figure S1**).

### Tumour-primed and tumour-resistant populations coexist in the basal layer

To gain further insights into the tumorigenic capacity of these distinct pools of progenitors, we used the potent oncogene BRAF^V600E/+^ (hereafter BRAF^V600E^), which encodes a different downstream oncogenic effector of the pro-proliferative MAPK signalling pathway. BRAF^V600E^ mimics the MAPK hyperactivation seen by the combination of BRAFi and *Hras* mutations. We generated a suite of mouse models expressing BRAF^V600E^ under the control of K14 and *Krt5* (K5-CreERT; hereafter K5) promoters targeting the stem-cell-like population and the IVL promoter targeting the committed basal populations (**Figures 2A and S2A**).

**Figure 2:**
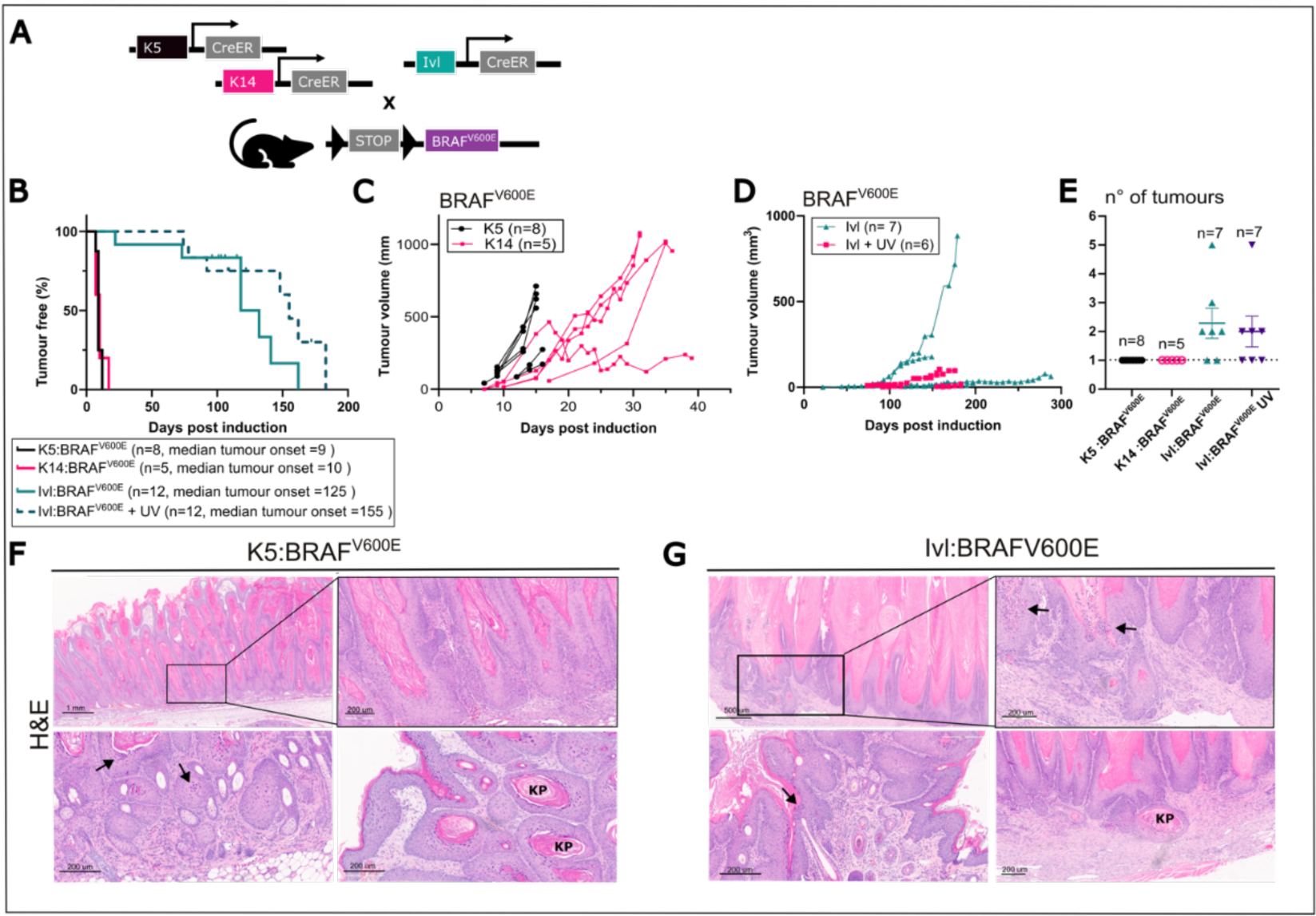
Tumour-primed and tumour-resistant populations coexist in the basal layer. (A) Schematic representation of the mouse models used for skin tumorigenesis. (B) Kaplan-Meier plot displaying tumour-free survival for K5:BRAF^V600E^, K14:BRAF^V600E^, Ivl:BRAF^V600E^, and Ivl:BRAF^V600E^ UV-treated female mice aged until clinical endpoint. (C-E) Total tumour growth in K5:BRAF^V600E^ and K14:BRAF^V600E^ (C) and in Ivl:BRAF^V600E^ and UV-treated Ivl:BRAF^V600E^ (D) female mice and total tumour number (Mean and SEM) (E) in female mice aged until clinical endpoint. (F-G) Representative H&E of K5:BRAF^V600E^ (F) and Ivl:BRAF^V600E^ (G) tumours at clinical endpoint showing keratin pearls are indicated (KP) and arrows indicate nuclear atypia.

Upon BRAF^V600E^ oncogene induction, both K5+ and K14+ cells rapidly formed tumours (9 and 10 days to tumour onset), without the need for further promotion, suggesting that these populations are permissive to tumour development and primed for oncogenic transformation (**Figures 2B and 2C**). In contrast, BRAF^V600E^ driven from the IVL model required a much longer time to develop tumours (125 days to tumour onset) and tumours grew slowly (**Figures 2B and 2D**). We also observed that the Ivl:BRAF^V600E^ model tends to develop multiple tumours at different body sites, including the back, paws, ears and lips, while the K5- and K14-driven BRAF^V600E^ models develop one tumour on their back at the precise area of tamoxifen application (**Figure 2E**).

Reducing the dose and duration of the tamoxifen-induction regimen in the K5:BRAF^V600E^ and K14:BRAF^V600E^ models delayed tumour onset, but tumours still grew rapidly (**Figure S2B**). Transplantation of subcutaneous pieces from K5:BRAF^V600E^-derived tumours into C57BL/6J and NSG-II2 immunodeficient mice revealed cancer stem cell potential and an ability to form secondary tumours **(Figure S2C)**. Thus, we termed this population tumour-primed.

The longer latency period for tumours to develop in the Ivl:BRAF^V600E^ model suggests that the IVL+ population requires additional cooperating events to facilitate tumour onset, so we termed this population tumour-resistant. To further promote cSCC in this population, we exposed the back of the Ivl:BRAF^V600E^ mice to mild UV [four weekly doses 6 standard erythema doses (SED)], but it did not accelerate tumour onset despite UV being a risk factor for cSCC due to its increase in mutation burden (**Figure 2B**). Similarly, topical treatment with the inflammation-promoting agent TPA (three times per week) did not accelerate tumorigenesis (**Figure S2D**). No statistical differences in tumour onset were seen between K14:BRAF^V600E^ cohorts housed in two different establishments or by sex (**Figures S2E and S2F**). However, the Ivl:BRAF^V600E^ model showed a slightly shorter tumour onset in the cohort housed at the CRUK SI compared to CRUK MI, while no differences were seen between sexes (**Figures 2G and 2H**).

We characterised the histopathological features of tumours originating in the K5:BRAF^V600E^, K14:BRAF^V600E^and Ivl:BRAF^V600E^ models and found remarkable similarities among them. All tumours were formed by vertical columns of keratinocytes with clearly demarcated limits (**Figure S3A**). Detailed immunohistochemistry showed distinctive pathological features of cSCC. These included enlarged interfollicular epidermis and cornified layers with an aberrant accumulation of extracellular keratin (keratin pearls), nuclear atypia, and incomplete maturation of keratinocytes (parakeratosis) as the keratinocytes in the cornified layer retain their nuclei (**Figures 2F, 2G and S3B**).

The tumours presented the hallmarks of papilloma with signs of squamous differentiation, as demonstrated by the increase in the number of layers, expressing the basal markers KRT5 and KRT14, but also the suprabasal marker KRT1 (**Figures S3C and S3D**). In the most advanced cases, these markers were expressed throughout the entire epidermal layer, indicating a complete loss of cellular hierarchy and disruption of multi-layer organisation. Moreover, the increase in Ki67 denotes epidermal hyperproliferation and hyperplasia of the interfollicular epidermis (**Figure S3E**). These lesions were highly proliferative, recapitulating the rapid onset and histopathology observed in human cSCC. Overall, despite both populations residing in the basal layer and sharing the microenvironment, they show profound differences in susceptibility to transformation upon oncogene induction.

### Oncogene-expressing tumour-resistant population takes over the entire skin epidermis but remains tumour-restrictive

We next aimed to understand how oncogene expression in the tumour-resistant population contributes to the clonal dynamics in the basal layer. In homeostasis, IVL+ marks a minority population in the basal layer that is committed to differentiation, although its differentiation does not prevent cell cycle entry ^21^. We followed in IVL+ or KRT14+ expressing populations by coexpressing the *Rosa26* LSL-tdTomato fluorescence protein (tdRFP) reporter in the presence and absence of BRAF^V600E^ (**Figure 3A**). Lineage-tracing experiments were conducted at different time points, ranging from before macroscopical changes appear in the skin up to day 160 or clinical endpoint, post tdRFP/oncogene induction (**Figures 3B**).

**Figure 3:**
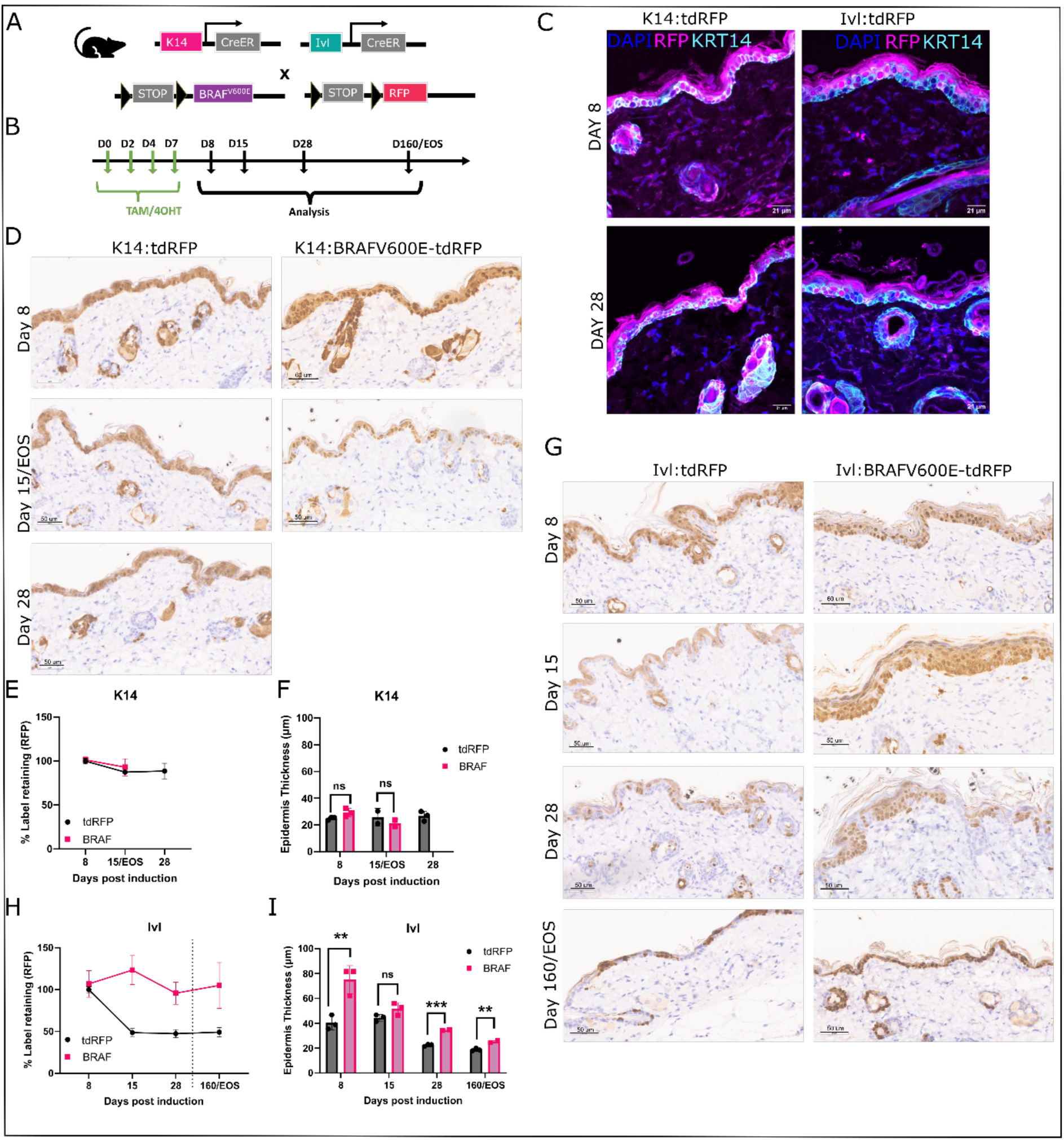
Oncogene-expressing tumour-resistant population takes over the entire skin but remains tumour-restrictive. (A) Schematic representation of the mouse models used for lineage tracing. (B) Schematic representation of the experimental design used for lineage tracing. (C) Representative Z-stack co-immunofluorescence (IF) of KRT14 and RFP at day 8 and day 28 post tdRFP induction in K14:tdRFP and Ivl:tdRFP normal skin. (D,G) Representative RFP IHC at different time points post-induction in K14:tdRFP and K14:BRAF^V600E^-tdRFP (D) and Ivl:tdRFP and Ivl:BRAF^V600E^-tdRFP (G). (E,H) Quantification of the clonal persistence in the basal epidermis region of K14:tdRFP and K14:BRAF^V600E^-tdRFP (E), and Ivl:tdRFP and Ivl:BRAF^V600E^-tdRFP (H) mice at different times post-induction. (F,I) Skin thickness quantification at different time points post-induction in K14:tdRFP and K14:BRAF^V600E^-tdRFP (F) and Ivl:tdRFP and Ivl:BRAF^V600E^-tdRFP (I). At least 10 measurements from three different skin stripes per animal. Points represent individual animals. Unpaired T-test. NS not significant P>0.05, ** p<0.01 and *** p<0.001.

First, co-immunofluorescence analysis with confocal microscopy of control skin (no oncogene) revealed that by day 8 in the K14:tdRFP model, all cells in the basal layer are labelled by RFP and remain labelled at day 28, marking a stable population as previously shown^19,21^. In the Ivl:tdRFP model at day 8, RFP labels only a small population of basal cells that co-expressed KRT14 and IVL transcripts, and some of those clones are maintained at day 28 (**Figure 3C**).

Then, we compared the RFP-labelled populations between control and oncogene-expressing skin. In the K14:tdRFP and K14:BRAF^V600E^-tdRFP models, all cells in the basal layer and their progeny in the suprabasal layer were labelled to saturation. Thus, there was no difference in clonal dynamics between groups (**Figure 3D**). An initial ∼10% drop in label-retaining cells was observed after day 8, when the induction regimen finished, but the population remained stable, renewing itself after that (**Figure 3E**). There were no differences in epidermis thickness between groups (**Figure 3F**).

In the Ivl:tdRFP model, the labelled cells were mainly located in the suprabasal layer with a few labelled cells in the basal layer at day 8 (**Figure 3G**). A ∼50% drop in the label-retaining population in the basal layer was seen after day 8, consistent with committed cell differentiating and delaminating (**Figure 3H**). Suprabasal cells that were not located under a label retainer IVL+ basal cluster turned over, and those regions completely lost the tdRFP label. Nevertheless, the labelled population remained stable for up to 160 days, marking a long-residing stable population of the basal layer able to self-renew and contribute with their progeny to the suprabasal layers (**Figures 3G and 3H**).

In contrast, upon oncogene activation in the Ivl:BRAF^V600E^ -tdRFP model, not only did we see a decrease in the label-retaining population after day 8, but we saw a ∼40% increase in the clonal persistence of labelled basal cells, which overtook the unlabelled population (**Figures 3G and 3H**). At day 15, the entire basal layer was labelled, similar to what we see in the K14:BRAF^V600E^tdRFP. Moreover, in the Ivl:BRAF^V600E^, there was a sharp increase in epidermal thickness, compared to control, due to the increased proliferation in differentiating cells that became smaller with time but remained significant during the study (**Figure 3I**). Thus, despite most basal cells carrying the oncogenic mutation, constraints remained in place resisting transformation.

These data show that oncogene expression in the committed IVL+ population led to a clonal proliferation dynamic similar to the stem-cell K14+ population but remained tumour-restrictive.

### cSCC shared transcriptional profile

Given the similarities in histology, we next conducted transcriptional profiling to identify potential vulnerabilities in these tumours. After batch correction to correct for any bias arising from batches sequenced at different times (**Figure S4A**), we merged the K5:BRAF^V600E^ and K14:BRAF^V600E^ tumour samples as they clustered independently of the model, and our analysis returned no differentially expressed genes (**Figures S4B and S4C**). Note that these tumours arose from the same cell populations simultaneously expressing KRT5 and KRT14. Then, we compared K14/K5:BRAF^V600E^- and Ivl:BRAF^V600E^-derived tumours against normal skin from control littermates. Unsupervised clustering and principal component analysis (PCA) showed that the tumours separated from the normal skin samples (**Figure 4A**).

**Figure 4:**
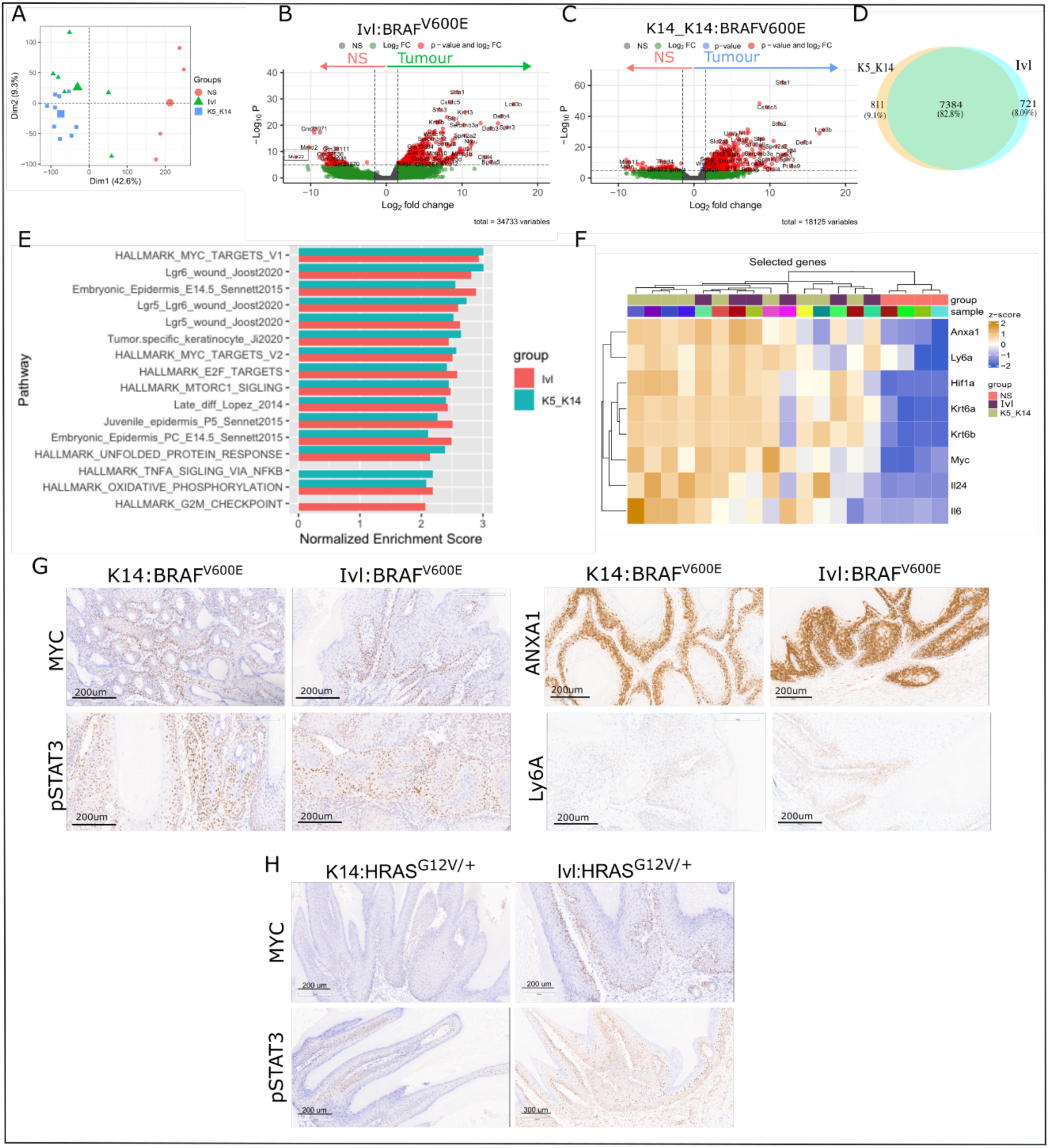
Shared transcriptional profile independent of the cell-of-origin or oncogene. (A) Principal component analysis (PCA) of normalised expression values showing the correlation between the transcriptional profiles of normal skin (NS, n=4), Ivl:BRAF^V600E^ (IVL, n=6) and K5/K14:BRAF^V600E^ (K5_K14, n=9). (B, C) Volcano plots showing differentially expressed genes in IVL tumours (B) and K5_K14 tumours (C) compared with normal skin using the Wald test (two-tailed). (D) Venn diagram showing upregulated genes shared between Ivl:BRAF^V600E^ (Ivl) and K5/K14:BRAF^V600E^ (K5_K14) tumours when compared with normal skin. (E) Gene set enrichment analysis of Hallmarks and indicated pathways in K5_K14 tumours and IVL tumours. Showing pathways significantly enriched (padj<0.01 and NES>2 based on an adaptive multi-level split Monte-Carlo). (F) Hierarchical clustering heatmap of selected genes showing normalised expression and correlation between normal skin (NS), Ivl:BRAF^V600E^ (Ivl) and K5/K14:BRAF^V600E^ (K5_K14). (G,H) Representative images of IHC validation of selected targets in K14:BRAF^V600E^ and Ivl:BRAF^V600E^ (G) and BRAFi treated K14:HRAS^G12D/+^ and Ivl:HRAS^G12D/+^ tumours (H) at clinical endpoint.

Both tumour groups revealed a large number of transcriptionally upregulated genes when compared to normal skin (**Figures 4B and 4C**). Nevertheless, the vast majority of upregulated genes (82.8%) were shared between the K14/K5:BRAF^V600E^ and Ivl:BRAF^V600E^ tumours (**Figure 4D**). Importantly, many transcriptional pathways we would expect to induce by promoters such as TPA are activated by the oncogene alone. Gene set enrichment analysis highlighted commonly upregulated signatures, including those associated with the *Myc*, *Lgr5* and *Lgr6* wound healing response ^23^, embryonic epidermis ^24^, and the human-derived tumour-specific keratinocyte population ^25^ among others (**Figure 4E and S5A**). *Myc* has been identified in wound healing as a crucial transcription factor during sebaceous duct dedifferentiation, generating a broad range of interfollicular epidermis cells that contribute to re-epithelialisation ^5^.

We also examined the expression of selected differentially expressed genes enriched in the tumours (**Figure 4F**). We observed high expression of *Myc*, the known cSCC markers *Krt6a* and *Krt6b* ^26^, and *Anxa1* ligand, which is secreted by wounded epithelia during regeneration and re-epithelisation ^23^. ANXA1 is also enriched in the tumour-specific keratinocyte population signature ^25^, and it has been shown to play a tumour-promoting role in other cancers of epithelial origin, such as cancers of the prostate ^27^ and breast ^28^. We also found increased expression of the basal markers *Ly6a* ^23^ and *Anxa1*, which have been proposed as a marker of foetal reprogramming and regenerative stem cells in the regenerating colonic epithelium and colorectal tumours (Gil Vazquez et al., 2022; Yui et al., 2018).

We also observed increased expression of the interleukins *Il24* and *Il6*. Interestingly, a recent model for tissue injury-sensing uncovered the role of IL24, produced at wound sites in response to hypoxia, in promoting epithelial proliferation and re-epithelialisation via pSTAT3 activation ^31^. This pathway may also extend to cSCC, as we see increased expression of the hypoxia sensor HIF1α and IL24 and pSTAT3 in our models (**Figure 4F)**. In addition, IL6 stimulates the release of proinflammatory cytokines from the cutaneous microenvironment in response to wounding, activating TGFβ receptor signalling and the STAT3 signal transduction pathway ^31,32^.

These transcriptomic results were validated by RNAscope for *Ly6a* and immunohistochemistry for MYC, ANXA1 and pSTAT3 in our models (**Figure 4G and 4H**). Similar findings were seen in tumours originating from the hair follicle stem cells upon MAPK signalling hyperactivation and deletion of the TGFβ receptor 1 (*Alk5*) (Lgr5:BRAF^V600E^-ALK5^fl/fl^, Lgr5:HRAS^G12V/G12V^-ALK5^fl/fl^, Lgr5:KRAS^G12D/+^-ALK5^fl/fl^)^10^ and deletion of *Trp53* and *Notch2* (Lgr5:TRP53^fl/fl^-NOTCH2^fl/fl^) (**Figure S5B**).

Overall, these results reveal shared molecular and transcriptional hallmarks of epidermal transformation independently of the cell-of-origin, tumour latency, or the oncogenic driver mutations.

### SOX2 is expressed in the tumour-resistant population

We next assessed whether transcriptional profiling could identify potential long-lasting markers of the cell-of-origin of these tumours. Thus, we directly compared the transcriptomic profile of tumours from K5/K14:BRAF^V600E^ and Ivl:BRAF^V600E^ model, which revealed only a handful of differentially expressed genes (**Figure 5A**). Notably, SOX2 emerged as a key transcriptional difference, being highly enriched in Ivl:BRAF^V600E^ tumours, relative to K5/K14:BRAF^V600E^.

**Figure 5:**
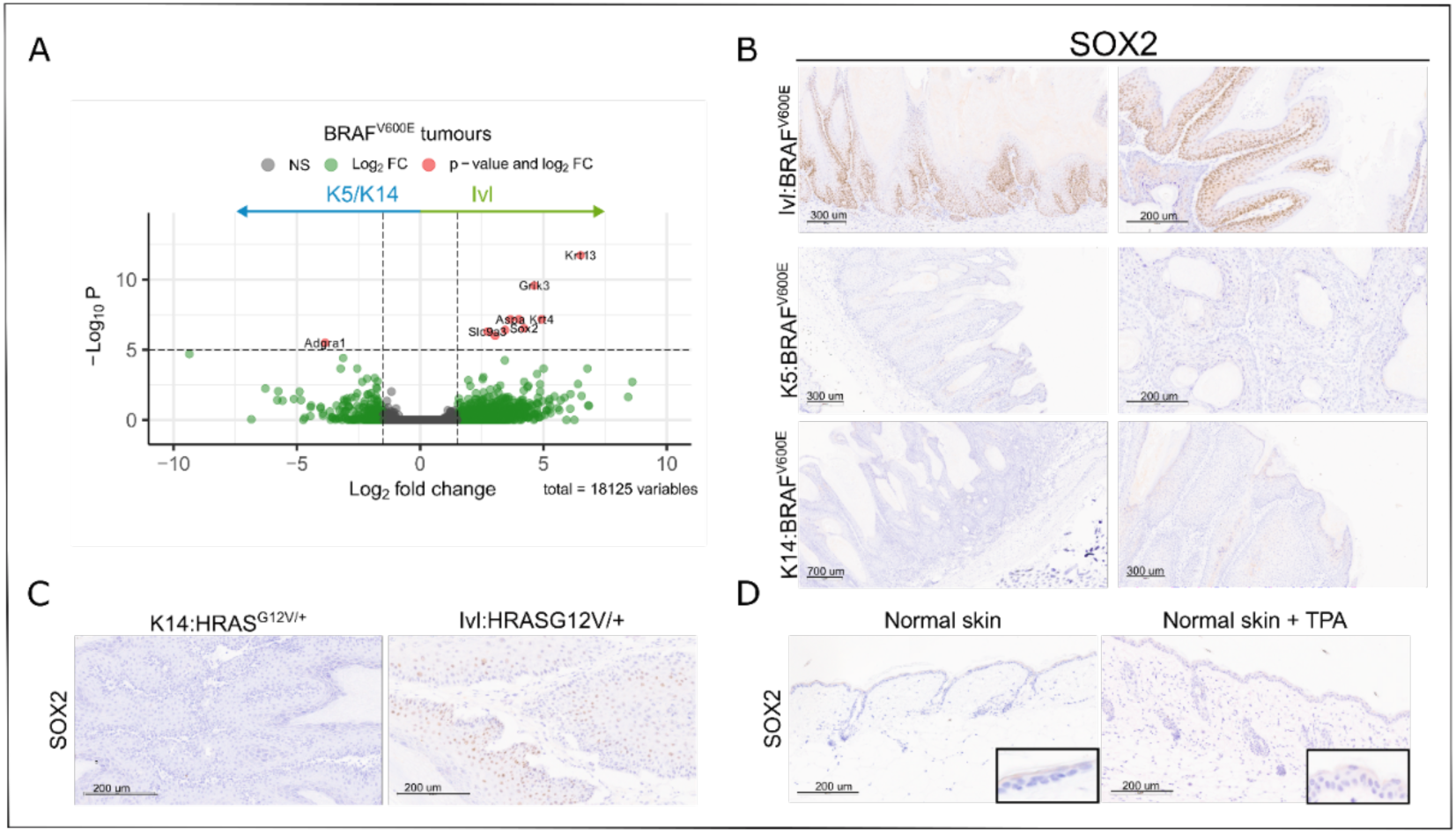
SOX2 is expressed in the tumour-resistant population. (A) Volcano plot showing differentially expressed genes between K5/K14:BRAF^V600E^ and Ivl:BRAF^V600E^ tumours using Wald test (two-tailed). (B,C) Representative images of IHC validation of SOX2 in Ivl:BRAF^V600E^, K5:BRAF^V600E^, and K14:BRAF^V600E^ (B) and BRAFi treated K14:HRAS^G12D/+^ and Ivl:HRAS^G12D/+^ tumours (C) at clinical endpoint. (D) Representative images of IHC of SOX2 in normal skin (left) and normal skin treated with TPA (right).

SOX2 has been identified as a super pioneer transcription factor able to bind to close chromatin through its ability to interfere with the maintenance of DNA methylation ^22^. Furthermore, SOX2 is commonly used as a marker of cancer stem cells and is upregulated in cSCC patients, where it rewires cells for tumour initiation and growth ^33,34^. We confirmed SOX2 enriched expression in Ivl:BRAF^V600E^ compared to K5:BRAF^V600E^, K14:BRAF^V600E^ profile by immunohistochemistry (**Figure 5B**). Similar results were obtained in HRAS-driven models where SOX2 expression was only seen in those arising from the IVL+ population (**Figure 5C**). Meanwhile, SOX2 expression was negligible in normal skin during homeostasis or in TPA-treated skin, suggesting that its expression is not linked to inflammation **(Figure 5D**). We then extended our investigation to a broader suite of models. We found SOX2 expressed in DMBA/TPA-induced tumours but not in tumours originating from *Lgr5+* cells (hair follicle bulge) driven by different oncogene combinations (**Figures S6A and S6B)**.

To highlight the cell-specific role of SOX2, we assessed the expression of SOX9, a different master regulator of chromatin accessibility that switches embryonic epidermal cells to a hair follicle stem cell fate and is constitutively active during tumorigenesis ^35^. As reported, we found that SOX9 was expressed mainly in the hair follicle cells in healthy skin homeostasis, and its expression was increased in the basal layer of all the tumours irrespective of the cell of origin (**Figure S6C**). The suprabasal markers KRT4 and KRT13, commonly expressed in skin homeostasis, were also differentially expressed between the K5/K14:BRAF^V600E^ and Ivl:BRAF^V600E^ models (**Figure 5A**). However, mRNA expression levels did not correlate with increased protein levels, as we found no significant differences in immunostaining intensity across our models (**Figure S6D**).

These data indicate that once the tumour-resistant population expresses SOX2, it drives and mimics the transcriptional profile of the tumour-prime stem population required cell status for tumorigenesis.

### SOX2 overexpression is sufficient to drive tumorigenesis from the tumour-resistant population

Given the potential aetiological activation of SOX2 in skin tumorigenesis, we assessed whether it is required for cSCC development in a cell-specific manner. We crossed the K14:BRAF^V600E^ and Ivl:BRAF^V600E^ models with mice harbouring a conditional *Sox2* knockout allele (SOX2^fl^) (**Figure 6A**). While conditional *Sox2* deletion did not affect tumorigenesis or tumour growth in the K14:BRAF^V600E-^ SOX2^fl/fl^ model (**Figure 6B**), it significantly delayed tumour onset and tumour growth in the Ivl:BRAF^V600E-^SOX2^fl/fl^ model (**Figure 6C**). Loss of *Sox2* had no impact on skin homeostasis in either population, consistent with its lack of expression in normal skin. Similarly, the deletion of *Sox2* had no effect on tumour growth in the K14 model but delayed tumour growth in the IVL model (**Figures 6D and 6E**)

**Figure 6:**
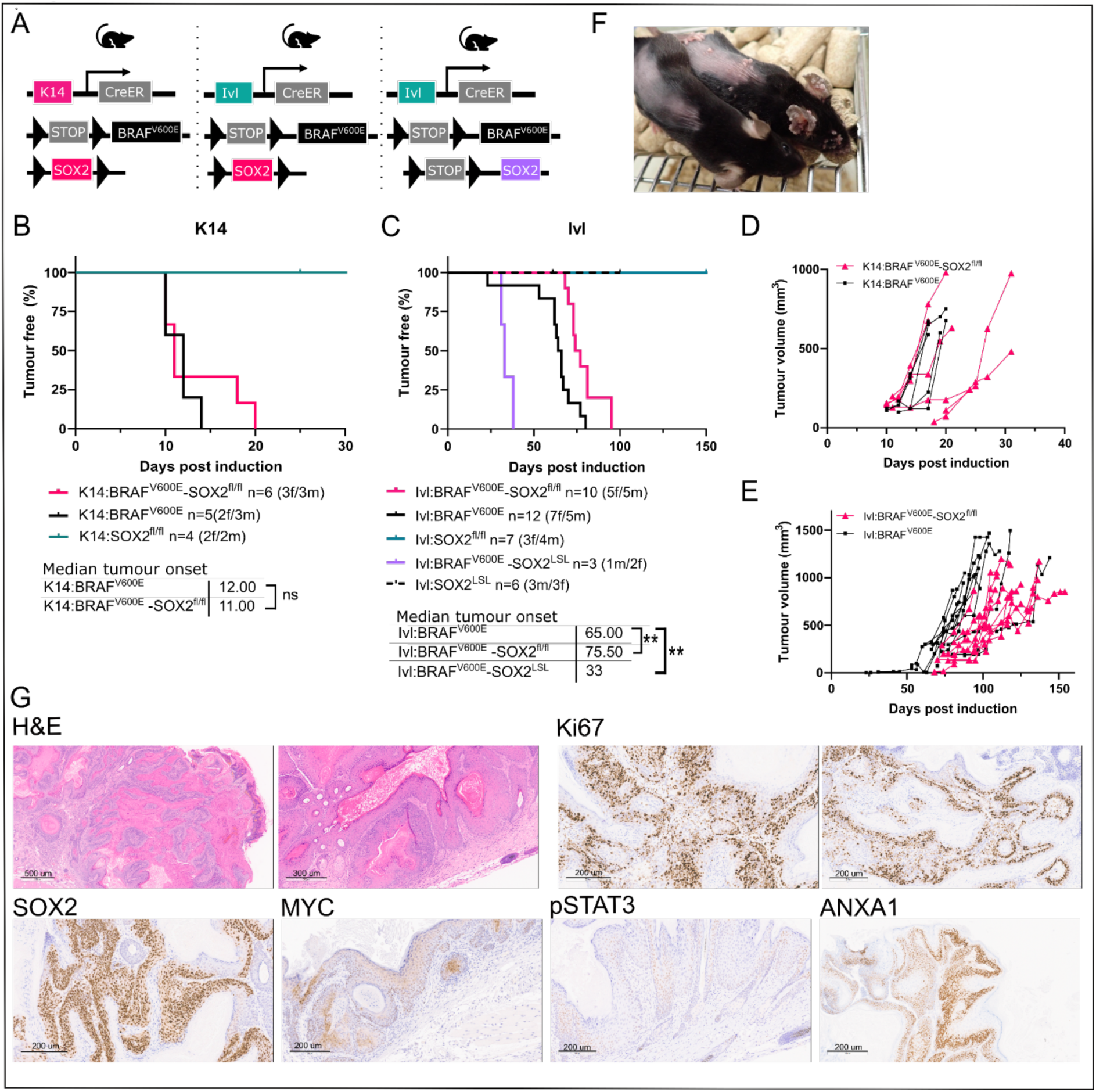
SOX2 overexpression is sufficient to initiate cSCC from the tumour-resistant population. (A) Schematic representation of the mouse models used for tumorigenesis. (B-E) Kaplan-Meier tumour-free survival plot and tumour growth for K14:BRAF^V600E^, K14:BRAF^V600E^-SOX2^fl/fl^ and K14:SOX2^fl/fl^ (B,D), and for Ivl:BRAF^V600E^, Ivl:BRAF^V600E^-SOX2^fl/fl^, Ivl:SOX2^fl/fl^, Ivl:BRAF^V600E^-SOX2^LSL^, and Ivl:SOX2^LSL^ (C,E). P-values were determined using the log-rank (Mantel-Cox) test. NS not significant P>0.05, ** p<0.01. Note that K14:BRAF^V600E^ and Ivl:BRAF^V600E^ cohorts correspond to CRUK-SI cohorts also shown in Figure S2 and are shown here for comparison. (G) Representative H&E and IHC of Ki67, SOX2, MYC, pSTAT3 and ANXA1 in Ivl:BRAF^V600E^-SOX2^LSL^ tumour at clinical endpoint.

The histopathological features of K14:BRAF^V600E-^SOX2^fl/fl^ and Ivl:BRAF^V600E-^ SOX2^fl/fl^ tumours were similar to those expressing the wild-type *Sox2* allele, including an enlarged epidermis displaying vertical columns of keratinocytes, incomplete maturation of keratinocytes, increased proliferation indicated by Ki67 immunoreactivity mainly in the basal layer, and MYC and pSTAT3 activation (**Figure S7A**). Interestingly, the rare Ivl:BRAF^V600E-^ SOX2^fl/fl^ tumours show patches escaping recombination and clones that retain SOX2 expression, as demonstrated by immunohistochemistry, which highlights its critical role in tumorigenesis (**Figure S7B**). SOX2+ escapee patches were not observed in the K14:BRAF^V600E-^SOX2^fl/fl^ tumours (**Figure S7B**).

Then, we assessed whether the gain of function of SOX2 could render IVL+ cells competent to tumorigenesis when coupled to a BRAF mutation. To this end, we interbred an allele which allows the inducible overexpression of SOX2 from the Rosa26 locus [*Rosa26*LSL-SOX2-IRES-eGFP (hereafter SOX2^LSL^)] to the tumour-resistant population in combination with BRAF^V600E^ in a tamoxifen-inducible manner. Upon SOX2 overexpression, the oncogene-targeted IVL+ population gave rise to fast-growing tumours. SOX2 significantly accelerated tumour onset from 65 to 33 days (**Figure 6F)**, while SOX2 overexpression on its own showed no phenotype (**Figure 6C**).

The Ivl:BRAF^V600E^-SOX2^LSL^ tumours show advanced histological features of cSCC, including poorly differentiated epithelial, keratin pearls, parakeratosis and invasion. These tumours show high and disorganised proliferation beyond the basal layer and activation of the previously defined shared transcriptional profile including pSTAT3, ANXA1 and MYC together with high expression of SOX2 (**Figure 6G**)

Overall, these data indicate that SOX2 is a cell-specific requirement for the IVL+ tumour-resistant population to acquire the competence to initiate cSCC. Thus, SOX2 gain of function in this population in combination with an oncogene leads to a clonal behaviour similar to the tumour-primed population and mediates the re-acquisition of stemness in the skin basal compartment.

### SOX2 drives developmental and juvenile epidermal programs and hints at the cell-of-origin

To determine whether SOX2 expression controls stemness in the skin in the absence of oncogene expression, we first immortalised two healthy human epidermal keratinocyte cell lines, derived from a juvenile and an adult donor, and subsequently overexpressed SOX2 in the derivative cell lines (**Figures 7A and S8A**). Using these human cell lines, we study the transcriptomic changes induced by SOX2 overexpression in homeostatic conditions.

**Figure 7:**
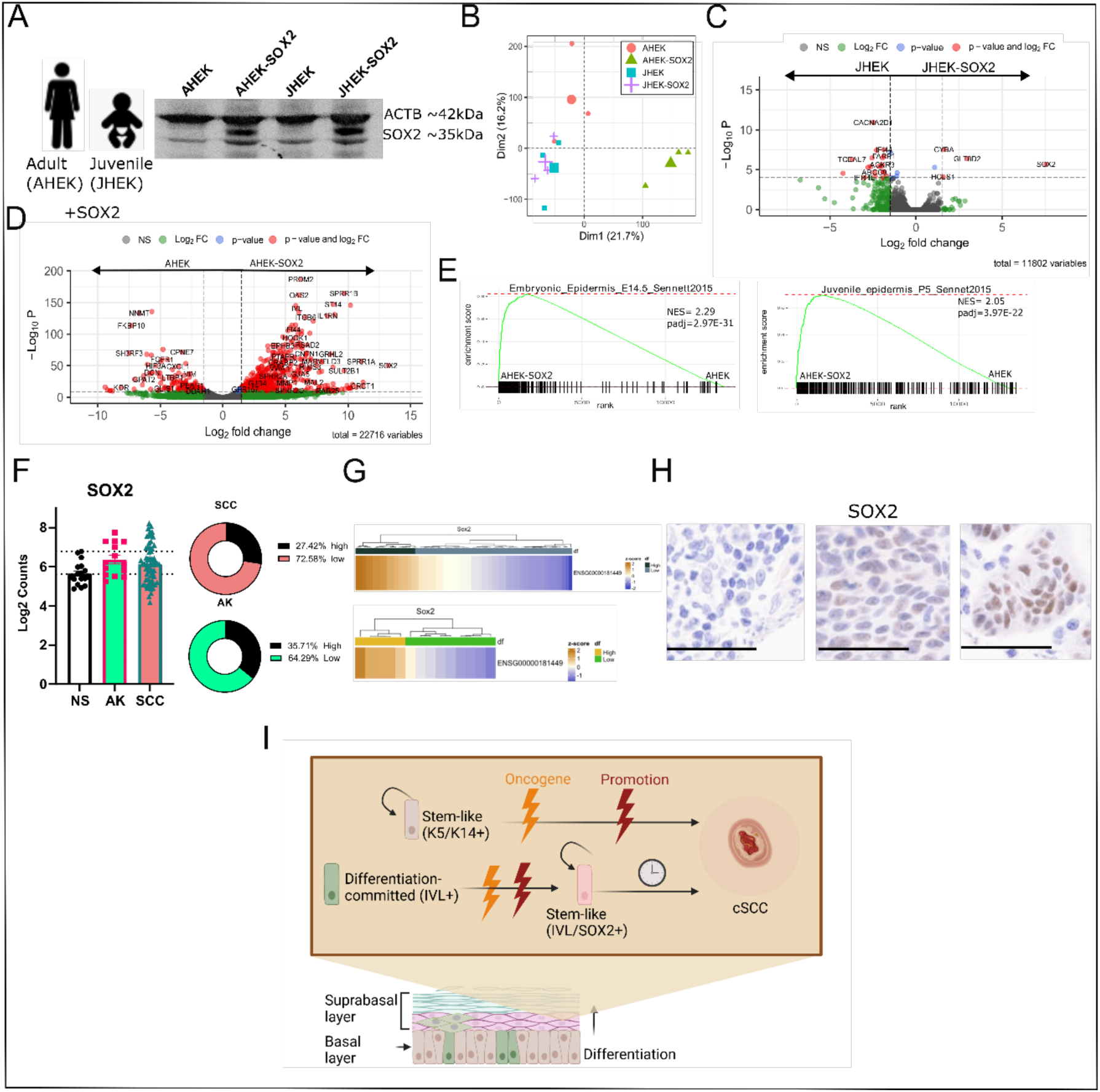
SOX2 drives developmental and juvenile epidermal signatures and hints at the cell-of-origin for cSCC patients. (A) Western-blot validation of SOX2 overexpression in adult (AHEK) and juvenile (JHEK) human keratinocytes. (B) Principal component analysis (PCA) of normalised expression values showing the correlation between the transcriptional profiles of JHEK, JHEK-SOX2, AHEK, and AHEK-SOX2. N=3, repeated independent experiments per group. (C,D) Volcano plot showing differentially expressed genes between JHEK and JHEK-SOX2 (C) and AHEK and AHEK-SOX2 (D) using the Wald test (two-tailed). (E) Gene set enrichment analysis of Embryonic epidermis and juvenile transcriptional signatures ^24^ in AHEK and AHEK-SOX2. Pathways are significantly enriched (padj<0.01 based on an adaptive multi-level split Monte-Carlo). (F) Normalised RNA expression levels (counts) of SOX2 from a cSCC human dataset ^17^ including normal skin (NS, n=17), and different levels of disease progression including actinic keratosis (AK, n=14) and cutaneous squamous cell carcinoma (SCC, n=66). Dotted lines mark the mean *SOX2* normalised expression in normal skin and two standard deviations from the mean. Pie charts showing the percentage of samples with high and low *SOX2* normalised expression for cSCC and AK groups. Two standard deviations from normal skin mean were used as a threshold. (G) Heatmap showing individual *SOX2* normalised expression levels in AK and cSCC groups. (H) Representative IHC of SOX2 conducted in human tissue microarray containing 250 cSCC histocores described in^36^. 196 (78.4%) of the samples were negative, and 54 (21.6%) were positive for SOX2 protein expression. The scale bar is 50 µm. (I) Graphic abstract of the model arising from this study. cSCC can be initiated from K14/K5+ stem-like cell or differentiation committed IVL+, the latter will require SOX2 activation and will take longer to escape constraints and trigger tumorigenesis. SOX2 reprogramming will activate embryonic and stem-like features critical for tumorigenesis. During this process, actively dividing IVL+ expand through the entire epithelium with little histological change, preparing for tumour initiation and progression and leading to an alternative route for tumorigenesis. We propose a common oncogenic programme, seen in human cSCC^25^ and inducible by inflammation, injury or paradoxical activation of MAPK, that allows the transformation of “normal” epithelial-containing oncogenes into a tumour without additional oncogenic events.

Interestingly, while SOX2 overexpression triggered transcriptomic changes in the adult cell line, separating it from the parental cell line, only small transcriptomic changes were seen in the juvenile SOX2-expressing cell line that clustered with its parental line (**Figures 7B-7D**). The profound SOX2-driven changes in the adult transcriptome included significant enrichment for the embryonic and juvenile epidermal signatures (**Figure 7E**). However, this enrichment was not observed in the SOX2 overexpressing juvenile cell line (**Figure 8B**). This could be due to the juvenile cell line resembling a stem-like population already expressing an embryonic/juvenile-like transcriptome, hence the lack of impact of SOX2 in these cells. These results resembled what we had seen in the models above; that the tumour-prime stem-cell population is not altered by the expression of SOX2, while in the differentiation committed tumour-resistant population, it drives the re-expression of stemness.

Finally, we sought to study how these data translate to human patients. We reanalysed a human dataset containing transcriptional data from different stages of disease progression ^17^. We observed that *SOX2* levels were increased in 27% of premalignant actinic keratosis (AK) and 35% of cSCC lesions, whereas *SOX2* expression levels were low in normal skin (NS) (**Figure 7F and 7G**). SOX2 levels did not correlate with the age of the patient or tumour aggressiveness (tumour depth or diameter) (**Figure S8C**). Furthermore, we divided these samples into SOX2 high and low groups and conducted further transcriptional comparisons, however, we did not see any major transcriptional changes either in the AK or cSCC groups (**Figures S8D and S8E**). Moreover, we looked at SOX2 protein expression in a tissue microarray containing 250 human cSCC^36^, which ranged from no expression, in the majority of the samples, to mid and high in 21.7% of the histocores (**Figure 7H**). Those results are in line with the ∼25% of human cSCC expressing high or medium levels of SOX2 reported previously ^33,34^.

This data aligns with the hypothesis that SOX2 is required to drive the re-expression of the stemness program needed to transform the tumour-resistant population, whereas it is not required for the transformation of the already stem-like basal population. Once SOX2 has rewired the epigenome and driven the embryonic and stem-cell pathways needed to trigger tumorigenesis, the tumours that result are indistinguishable from those originating from a stem population (**Figure 7I**).

These results underscore a diverse cell origin for cSCC, where SOX2 marks a subset of tumours emerging from the committed tumour-resistant population.

## Discussion

The heterogeneity of the interfollicular epidermis basal layer is well established. However, considerable debate remains as to whether it comprises two distinct progenitors or two populations that relate to different levels of commitment towards differentiation^19–21,37^. Our research uncovers the mechanism underlying the different tumour-initiating capacities following oncogenic mutations of both populations and identifies SOX2 as a critical enabler that renders committed progenitors susceptible to cSCC initiation.

Importantly, we generated novel genetically engineered mouse models that resemble the rapid cSCC development seen in melanoma patients treated with BRAFi, due to paradoxical MAPK signalling activation ^13–15^. BRAFi promoted the development of tumours from cells carrying HRAS^G12V^ mutations that had been sitting in the skin for up to 180 days without driving transformation. This shows that the skin can tolerate mutations in oncogenic pathways with little histological change but can rapidly progress to tumorigenesis given a specific stimulus or promoter.

We uncovered the mechanisms driving the difference in oncogene permissiveness by modelling MAPK signalling hyperactivation in the stem-cell K14/K5+ population and the committed progenitor IVL+ population. In particular, we obtained rapidly developing tumours within days of oncogene induction in the K14/K5+; hence, we called this population tumour-primed. However, oncogene induction in the IVL+ differentiation committed basal population resulted in homeostasis and normal-looking skin despite the oncogene-bearing cells colonising the entire epithelium within a week. Thus, we termed this population tumour-resistant. Expression of HRAS^G12V^ or KRAS^G12D^ oncogenes in the skin epithelium also outcompeted the wild-type population in homeostasis ^38^. Furthermore, oncogene expression in the IVL+ cells increased their residence in the basal layer, preventing delamination and allowing for a longer latency in the progression to tumorigenesis. Once tumours developed, they resembled those originating from K5+/K14+ cells.

Transcriptional profiling of the tumours arising from an IVL+ or a K5+/K14+ revealed a common transcriptional landscape that is enriched for an activated wound-healing response ^23^, embryonic developmental pathways ^24^ and a tumour-specific keratinocyte population seen in human cSCC ^25^. In addition, we observed strong MYC activation across all tumours, which has been proposed as a mechanism to confer stem cell properties and dedifferentiation in the *Gata6*+ epidermal populations^5^. At the same time, we revealed evidence of hijacking the tissue injury-sensing and repair pathway, triggered by hypoxia and driven by IL24 and STAT3 activation that resolves in a coordinated tissue repair after wounding ^31^. Similar to wounds, the core of a tumour is hypoxic, and we observed increased transcription of *Hif1a* and *Il24*, as well as STAT3 phosphorylation that enhanced oncogene-driven proliferation to generate the critical tumour mass required for progression. These results highlight the close interplay between proliferation during tumorigenesis and re-epithelisation during wound sealing.

Much of our understanding of cSCC in the mouse has come from seminal work using DMBA/TPA^2,11^ or more lately mouse models that have focussed on using stem cell /progenitor to drive KRAS or other MAPK mutations in combination with TP53^8,10,17^. These models typically result in widespread oncogenic mutations across the skin, yet only a single tumour often emerges, mirroring human studies that highlight the skin’s ability to tolerate oncogenic mutations. Despite the diversity of oncogenic events across these models, they converge on a core set of skin transformation pathways, as shown here. These pathways can be rapidly triggered by BRAF mutation or by paradoxical MAPK activation via BRAFi in HRAS-mutant cells, leading to tumorigenesis within days or over months to years in models with longer latency. We propose that the acquisition of this oncogenic programme —also seen in human cSCC^25^ and inducible by inflammation, injury or paradoxical activation of MAPK— allows the transformation of “normal” epithelial-containing oncogenes into a tumour without requiring additional oncogenic events.

One of our key findings was that SOX2 activation in the tumour-resistant population is sufficient to render this population susceptible to transformation. Knocking out this gene *in vivo* delayed tumour onset and growth but did not affect the tumour-primed population. Conversely, SOX2 overexpression in combination with oncogene expression rapidly accelerated tumorigenesis *in vivo,* showing that SOX2 is both necessary and sufficient for transforming the IVL+ lineage.

The limited capacity of the IVL+ basal population to switch to symmetric division and its tendency to preferentially undergo asymmetric division is part of its transcriptome-engrained commitment towards differentiation^19,21^ and could explain its tumour-resistant phenotype also seen in basal cell carcinoma^37^. Indeed, in basal cell carcinoma, only the stem-cell population and not the progenitors have been proposed to initiate tumorigenesis^37^. However, we argue that both populations (stem-like and committed) can trigger skin tumorigenesis.

There is evidence that SOX2 activation happens in ∼20% of cSCC patients^33,34^. This suggests that the IVL+ basal population acts as a cell-of-origin for these tumours and that SOX2 may serve as a marker for classifying human cSCC according to the cell of origin. Still, the committed progenitors are tumour-resistant, and they need to re-acquire the stemness features to stop differentiation and apoptosis and allow transformation. Thus, this requirement of a stemness transcriptome, acquired by SURVIVIN expression in basal cell carcinoma^37^ or by SOX2 in cSCC reconciliates our results. The convergent “stemness” state required for tumour initiation is also found in tumours developed from the DMBA/TPA model ^2^.

Although the role of SOX2 in cSCC initiation has been previously described^33,34^, our study shows that its expression is only required and essential in the tumour-resistant population, where it confers the stemness required to overcome their inability to initiate cSCC. SOX2 is a super pioneer factor able to bind closed chromatin and rewire the epigenome to drive stem cell features ^22,39^. Indeed, we show that SOX2 overexpression in homeostasis can drive embryonic and juvenile transcriptomes. Thus, SOX2 would allow the tumour-resistant population to suppress, freeze, or completely abolish its commitment towards differentiation by activating the stemness profile required for transformation. IVL+ cells may dedifferentiate and become IVL- to drive tumorigenesis and to remain in the basal layer, where the proliferative environment is confined. Indeed, cancer stemness is not restricted to the unidirectional route (stem cell to progenitors) that applies in normal tissues, as shown by therapeutic agents inducing different differentiation states^2^.

We hypothesise that the more fluid and gradual differentiation mechanism recently proposed^21^ could be reverted towards dedifferentiation as the committed cells residing in the basal layer have not necessarily exited the cell cycle. This model could explain the almost identical transcriptomic profile obtained from tumours derived from the K14/K5+ population and the Ivl+ population, as both populations will go on to delaminate, which argues against the presence of two distinct epidermal populations.

How SOX2 is activated by oncogenic mutations is still elusive. It may be directly or indirectly downstream of the commonly activated pERK or MYC signalling pathways. A deep understanding not only of the tissue-specific mechanisms that drive transformation but also of the cell-specific vulnerabilities that govern epithelial maintenance in homeostasis and tumorigenesis will allow the development of rational cancer therapies to contain tumours at their source.

Collectively, we have generated new genetically driven *in vivo* models to study cSCC and identified SOX2 as a key regulator of the different susceptibility of the tumour-primed and tumour-resistant skin populations. SOX2 expression in oncogene-carrying tumour-resistant clones increases their residence time in the basal layer and prevents delamination by driving a stem-like transcriptomic profile that enables this population to initiate tumorigenesis at a longer latency.

## Online Methods

### Genetically engineered mouse models and husbandry

Mice were housed in accordance with UK Home Office Regulations, maintained in a pathogen-free facility under a 12-h light-dark cycle at a constant temperature between 19–23°C, and 55 ± 10% humidity, and given drinking water and fed standard chow diet *ad libitum*. Experiments conducted at the CRUK Scotland Institute Animal Facility were in accordance with the UK Home Office guidelines, under project licence PP3908577, and were reviewed and approved by the University of Glasgow Animal Welfare and Ethical Review Board (AWERB). Experiments conducted at the CRUK Manchester Institute were performed at Alderley Park Animal Research Unit, with breeding at the University of Manchester incubator breeding facility, in accordance with UK Home Office regulations, under project licences PE4369EDB and P671A5B06, and reviewed and approved by the CRUK Manchester Institute’s AWERB. Animals were monitored regularly until terminally sacrificed by schedule 1 at clinical endpoint: either when the cumulative tumour burden had reached a maximum volume of 1500 mm^3^ or when a single tumour exceeded 15 mm in diameter, weight loss >20%, or any other signs of ill health and distress. For all mouse studies, no formal randomisation was performed, and researchers were not blinded to the mouse genotypes. No exclusions were performed and all mice included in experimental cohorts were included in the analysis.

Genotyping was performed by Transnetyx according to the previously published protocols provided in the references for each allele. Mice were maintained on a C57BL/6J background, carrying the following alleles/transgenes: *Ivl*-CreERT2 ^8^, *Krt5*-CreERT2 ^40^, *Krt14*-CreERT2 ^41^, BRAF^V600E^ ^42,43^, HRAS^G12V^ ^44^, *Rosa26LSL-tdRFP* ^45^, *Sox2^fl^* ^46^ and *Rosa26LSL-SOX2-IRES-eGFP* ^47^. Archived tumour blocks [from^10^] included the additional following alleles: *Lgr5*-CreERT2 ^48^, *Kras*^G12D^ ^49^, *Notch2*^fl^ ^50^, and *Trp53*^fl^ ^51^.

Genetic recombination was induced in mice of either sex 8-12 weeks of age by topical application to the shaved backs of the mice of 100 µL tamoxifen (10 mg/ml in ethanol; T5648, Sigma) in models driven by Ivl-CreERT2, or 4 µl 4-hydroxytamoxifen (20 mg/ml in ethanol; H6278, Sigma) in models driven by *Krt5*-CreERT2 and *Krt14*-CreERT, unless otherwise stated. This was repeated three more times over 8 days.

### UV irradiation

UV irradiation experiments were conducted at the CRUK Manchester Institute husbandry facility following the UK Home Office regulations under project licences PE4369EDB and P671A5B06. In cohorts exposed to UVR, mice were subjected to UVR exposure once per week for 4 weeks, beginning 4 weeks after the final tamoxifen application. For this process, mice were anaesthetised by intraperitoneal injection with 1 mg/kg Domitor and 100 mg/kg ketamine, with 5 mg/kg Antisedan anaesthetic reversal. The backs were shaved, and a black cloth was used to cover regions that were to remain unexposed. The Waldmann UV181 unit with UV6 broad wavelength (280-380 nm) lamp was used for UVR exposure and the intensity was tested regularly with a USB2000+ spectroradiometer (Ocean Optics). The UV dose used was 0.6 kJ/m^2^, which equated to 3 minutes and 14 seconds of exposure. Following irradiation, E45 moisturising cream was topically applied to the back.

### Tumour fragment implantation

Tumour implantation experiments were conducted at the CRUK Manchester Institute husbandry facility following the UK Home Office regulations under project licences PE4369EDB and P671A5B06. A tumour derived from a K5:BRAF^V600E^ mouse was resected upon clinical endpoint under a laminar flow hood aseptically. The tumour was cut into 2-3 mm^3^ pieces for immediate implantation on anesthetised 6-8 week-old C57/6J littermates and NSGII2 mice. Briefly, recipient animals were anesthetised with isoflurane and kept on a heated stage. A tumour piece was implanted subcutaneously between the skin and the peritoneal wall. The surgical incision was closed using a surgical clip. Rymadyl/Carprofen analgesic was given at 4mg/kg. Mice were monitored until consciousness was regained and then monitored daily. The surgical clip was removed 7 days post-surgery.

### *In vivo* drug treatments

*In vivo* drug treatments were conducted at the CRUK Scotland Institute Animal Facility following the UK Home Office guidelines, under project licence PP3908577. Skin inflammation was induced by topically applying 150 µl of TPA (31.25 µg/ml) in acetone (Sigma-Aldrich 16561-29-8) to shaved dorsal skin three times per week until tumour signs appeared. The BRAFi Dabrafenib was administered by daily oral gavage (30 mg/kg in 100 µl) during the experiment.

### Immunohistochemistry and immunofluorescence

Organs collected in 10% neutral buffered formalin were stored at room temperature for 20-28 hours, followed by transfer to 70% ethanol and storage at 4°C until processing. All haematoxylin and eosin (H&E), immunohistochemistry, co-immunofluorescence and *in situ* hybridisation staining were performed on 4-µm formalin-fixed paraffin-embedded sections (FFPE) which had previously been heated at 60°C for 2 hours.

The following antibodies were used on a Leica Bond Rx autostainer: KRT5 (905501, Biolegend), Ki67 (12202, Cell Signaling), SOX2 (14962, Cell Signaling) and pSTAT3 (9131, Cell Signaling). All FFPE sections underwent onboard dewaxing (AR9222, Leica) and epitope retrieval using ER2 solution (AR9640, Leica) for 20 minutes at 95°C. Sections were rinsed with Leica wash buffer (AR9590, Leica) before peroxidase block was performed using an Intense R kit (DS9263, Leica) for 5 minutes. Sections were rinsed with wash buffer before primary antibody application at an optimal dilution (KRT5, 1/1500; Ki67, 1/1000; pSTAT3, 1/100; SOX2, 1/200). The sections were rinsed with wash buffer before the application of anti-rabbit EnVision HRP-conjugated secondary antibody (K4003, Agilent) for 30 minutes. The sections were rinsed with wash buffer, visualised using DAB, and counterstained with haematoxylin from the Intense R kit.

FFPE sections for KRT14 (ab7800, Abcam), c-MYC (ab32072, Abcam), RFP (600-401-379, Rockland) and SOX9 (AB5535, Millipore) staining were stained on an Agilent autostainer Link48. FFPE sections were loaded into an Agilent pre-treatment module to be dewaxed and undergo heat-induced epitope retrieval (HIER) using a High pH target retrieval solution (K8004, Agilent). After HIER, the sections were rinsed in FLEX wash buffer (K8007, Agilent) before being loaded onto the Agilent autostainer. The sections underwent peroxidase blocking (S2023, Agilent) for 5 minutes and were rinsed with FLEX buffer. The primary antibody application was at an optimised dilution (KRT14, 1/300; c-MYC, 1/800; RFP, 1/1000; SOX9, 1/500). Sections were washed with FLEX buffer before application of anti-rabbit EnVision secondary antibody for 30 minutes. Sections were rinsed with FLEX wash buffer before applying Liquid DAB (K3468, Agilent) for 10 minutes. Sections were washed in water and counterstained with haematoxylin ‘Z’ (RBA-4201-00A, CellPath).

In-situ hybridisation detection for *Anxa1* (509298), *Ivl* (422538), *Ly6a* (427578), PPIB (313918; positive control) and dapB (312038; negative control) (all Bio-Techne) mRNA was performed using RNAScope 2.5 LSx (Brown) detection kit (322700; Bio-Techne) according to the manufacturer’s instructions. H&E staining was performed on a Leica autostainer (ST5020). Sections were dewaxed in xylene, taken through graded ethanol solutions and stained with haematoxylin ‘Z’ (RBA-4201-00A, CellPath) for 13 minutes. Sections were washed in water, differentiated in 1% acid alcohol, washed and the nuclei stained blue in Scott’s tap water substitute (in-house). After washing with tap water, sections were placed in Putt’s eosin (in-house) for 3 minutes. To complete H&E, IHC & ISH staining, sections were rinsed in tap water, dehydrated through a series of graded alcohols, and placed in xylene. The stained sections were coverslipped in xylene using DPX mountant (SEA-1300-00A, CellPath).

Sections for KRT14 and RFP co-IF staining were loaded onto a Leica Bond Rx autostainer. The FFPE sections underwent on-board dewaxing and epitope retrieval using ER2 solution for 20 minutes at 95°C. Sections were rinsed with Leica wash buffer (AR9590, Leica) before application of 10% normal goat serum (X090710-8, Agilent) for 30 minutes. The sections were rinsed with Leica wash buffer before application of anti-RFP antibody at 1/1000 dilution for 1 hour. Sections were rinsed with Leica wash buffer and goat anti-rabbit IgG 488 (A11034, Invitrogen) secondary antibody diluted 1/250 for 30 minutes. After rinsing with Leica wash buffer KRT14 antibody was applied at 1/750 dilution for 1 hour. Sections were rinsed with Leica wash buffer and goat anti-mouse IgG 647 secondary antibody (A21236, Invitrogen diluted) 1/250 for 30 minutes before application of DAPI (MBD0015, Sigma-Aldrich). To complete the staining, sections were mounted using ProLong Diamond antifade mountant (P36970, Thermo Fisher Scientific).

Immunohistochemistry images were acquired using a SCN400F slide scanner (Leica Microsystems) at ×20 or x40 magnification. Confocal images were collected on a Zeiss 710 point-scanning confocal microscope, built on an inverted Zeiss Axio Imager.Z2 stand. Images were acquired using a EC Plan-Neofluar 40x/1.30 Oil and a confocal pinhole diameter of 107 µm. Multi-channel images were captured sequentially: DAPI (nuclear marker) using 405 nm excitation and 410-481 nm emission bandwidth, IgG 488 using 488 nm excitation and 514-582 nm emission and IgG 647 using 633 nm excitation and 638-747 nm emission. Images were collected with a 1x zoom with an image size of 2320 x 2320 pixels, yielding a pixel size of 92 x 92 nm, and a 1.38 us pixel dwell time. Z-stacks were collected using a step size of 2 um. Images were acquired using the software Zen LSM 2.1 Black (Zeiss).

### RNAseq sequencing and analysis

RNA was extracted from fresh frozen tumour samples or human cell line pellets using an AllPrep DNA/RNA kit (74104, Qiagen) according to the manufacturer’s instructions. Indexed poly(A) libraries were prepared using 200 ng of total RNA and 14 cycles of amplification with the Agilent SureSelect Strand-Specific RNA Library Preparation Kit for Illumina Sequencing (G9691B, Agilent). RNA polyA libraries were sequenced using NovaSeq 6000 XP SP, 200 cycles, paired-end reads (2 x 101 bp) 20-30 million reads. RNA preparation and sequencing of mouse samples were performed at the CRUK Manchester Institute histology core and sequencing facility. RNA preparation and sequencing of human cell lines were performed at GENEWIZ Azenta.

For mouse samples raw sequence quality control was performed using FastQC (v0.11.8) before and after removing adapters and low-quality base calls (Phred score <20) using TrimGalore with default options (v0.6.6). Trimmed reads were aligned to GRCm38, release 100 using Hisat2 (v2.1.0). Gene counts were subsequently estimated using featurecounts (subread/1.6.3). For human cell lines, raw sequence quality control and alignment were performed using Nextfow nf-core/rnaseq pipeline using STAR and Salmon ^52^. Trimmed reads were aligned to GRCh38, release 113. Sva::ComBat-seq was used for batch correction (v3.52.0). For all samples, after removing transcripts without a minimum of 5 reads in at least one sample, the differential expression analysis between mouse tumours and skin was performed using the R package DESeq2 (v1.44.0). The resultant p-values were corrected for multiple comparisons using the Benjamini-Hochberg approach. The following additional R packages were used for downstream analysis: pheatmap (v1.0.8), fgsea (v1.30.0), enhancedVolcano (v1.22.0), clusterProfiler (v4.12.6), dplyr (v1.1.4), tidyr (v1.3.1), AnnotationHub (v3.12.0), ggvenn (0.1.10), AnnotationDbi (v1.66.0), mixOmics (v6.26.0).

### Culture of human keratinocyte cell lines and introduction of genetic modifications

Human keratinocyte cell lines were maintained in serum-free Keratinocyte Growth Medium 2 (C-20011, PromoCell) until confluent and then split using 0.25% Trypsin-EDTA (T4049-100ML, Merck) and quenched using Defined Trypsin Inhibitor (R007100, Gibco). All cell culture incubators were set to 37°C and 5% CO2.

Normal human primary juvenile keratinocytes (JHEK) (C-12006, PromoCell) and normal human primary adult keratinocytes (AHEK) (102-05a, Cell Applications) were immortalised with in-house produced lentivirus expressing the hTERT-hygromycin resistant pLV-EF1a-IRES-Hygro plasmid and selected with hygromycin (300 µg/ml). The plasmid was a gift from Tobias Meyer (85140, Addgene) ^53^). Then, nucleofection-based CRISPR-Cas-9 methodology was used to generate p16 KO using the 4D Nucleofector Core Unit (AAF-1002B with AAF-1002X, Lonza). A gRNA with enhanced stability targeting *Cdkn2a* (p16) was designed and purchased from Integrated DNA Technologies (IDT). Ribonucleoprotein (RNP) complexes were formed by diluting 120 pmol sgRNA, 104 pmol Cas9 Nuclease V3 (1081059, IDT), 1 µl of Alt-R Cas9 Electroporation Enhancer (1075916, IDT) in Nuclease-Free IDTE, pH 7.5 (1X TE solution) (I11-01-02-02, IDT). RNP complex was added to 20 µl of Primary Cell P3 Nucleofector Solution (V4XP-3032, Lonza) and nucleofected using programme CM137. The mixture was then transferred to a 20 µl nucleocuvette strip and nucleofected using programme CM137. The targeted sequence was PCR-amplified, sequenced at Eurofins and KO-validated using the TIDE algorithm (achieving >90% efficiency)^54^. *SOX2* overexpression lentivirus with puromycin resistance was purchased from VectorBuilder ready-to-use (vector id VB900088-3008dcy). SOX2-overexpressing keratinocytes were selected with 1 µg/ml puromycin. GFP overexpression control lentivirus with puromycin resistance purchased from VectorBuilder ready-to-use were used as controls. Cell pellets were regularly collected and submitted for cell line authentication at Eurofins and tested for Mycoplasma in-house.

### Immunoblotting

Proteins were extracted from whole-cell lysates using RIPA buffer (R0278, Sigma-Aldrich) mixed with 1 tablet of cOmplete™ Protease Inhibitor Cocktail (11873580001, Sigma-Aldrich). Cell suspensions were incubated on ice for 10 minutes before centrifuging at 13,000 rpm for 10 minutes at 4°C. Protein extracts from the supernatants were quantified using the Pierce™ BCA Protein Assay Kit (23225, Invitrogen). Protein extracts (25 μg) were denatured in 4x Laemmli sample buffer (1610747, BIO-RAD) at 95 °C for 5 minutes. The samples were separated on NuPAGE™ 4-12% Bis-Tris Mini Protein Gels (NP0322BOX, Invitrogen) at 120 V for 1 hour in 1x MOPS SDS running buffer (B0001, Invitrogen) together with the Chameleon Duo pre-stained Protein Ladder (928-60000, LI-COR). The proteins were transferred onto a nitrocellulose membrane (170-4270, BIO-RAD), using the BIO-RAD Trans-Blot Turbo Transfer System. The membrane was blocked with 1x TBST with 5% BSA (A7906, Sigma-Aldrich) for 1 hour at room temperature in the dark. The membrane was then incubated in anti-SOX2 antibody 1:1000 (23064, Cell Signalling) and anti-β-actin 1:10,000 (MA1-140, Invitrogen) diluted in 1x TBST 5% BSA, at 4°C overnight in the dark. β-actin was used as a loading control. Followed by 3x10 minute washes in 1x TBST in the dark and subsequently incubated in fluorescence-conjugated secondary antibodies, diluted in 5% BSA in 1x TBST, at room temperature for 45 minutes in the dark. Secondary antibodies used were anti-rabbit IgG (A32735, Thermo Fisher Scientific) and anti-mouse IgG (A21057, Thermo Fisher Scientific). This was followed by 3x10 minute washes in 1x TBST before the membrane was scanned using the LI-COR Odyssey® CLx Infrared Imaging System.

## Data availability

The data discussed in this publication have been deposited in NCBI’s Gene Expression Omnibus. The accession numbers for the RNA-seq data reported in this paper are NCBI GEO GSE280236 (mouse data) and NCBI GEO GSE281000 (human cell lines data). No new code was generated in this publication.

## Acknowledgements

We thank the Core Services and Advanced Technologies at the Cancer Research UK Scotland and Manchester Institutes, particularly the Biological Services Unit, Histology Service and Molecular Technologies. We also thank Claire Mitchell and the BAIR Biological Advance Image Resource service at Cancer Research UK Scotland Institute. We thank Kevin Haigis (Dana-Farber Cancer Institute) for providing the HRAS^G12V^ mouse model. We are grateful to Sarah Ressel and Shan Quah for the critical review of the manuscript and to Nathalie Sphyris for expert writing advice (Cancer Research UK Scotland Institute). This work was supported by Cancer Research UK Cancer Grand Challenges and The Mark Foundation for Cancer Research to the SPECIFICANCER team (A29055) and by CRUK core funding to the Cancer Research UK Manchester Institute (C5759/A27412) and the Cancer Research UK Scotland Institute (C5759/A31287). P.P.C. and C.C. were funded by the Cancer Research UK Grand Challenge SPECIFICANCER Consortium (A29055 & A27412). C.A.F, R.A.R., P.C., T.J, O.J.S. were supported by CRUK Beatson Institute core funding (A17196, A31287).

## Author Contributions

P.P.C., A.C., R.M. and O.J.S. designed experiments and interpreted results. P.PC, C.C., C.A.F., R.A.R., T.J.S. and P.C. performed experiments and analysed results. P.P.C. analysed publicly available human cancer data sets. P.P.C. processed and analysed the RNA sequencing data. P.PC., A.C. and O.J.S. wrote the paper, and reviewed and discussed the drafted manuscript. All authors contributed to the manuscript.

