## Supplementary Figures for "SOX2 empowers a rapid tumorigenic programme from the tumour-resistant population in the skin"

Figure S1

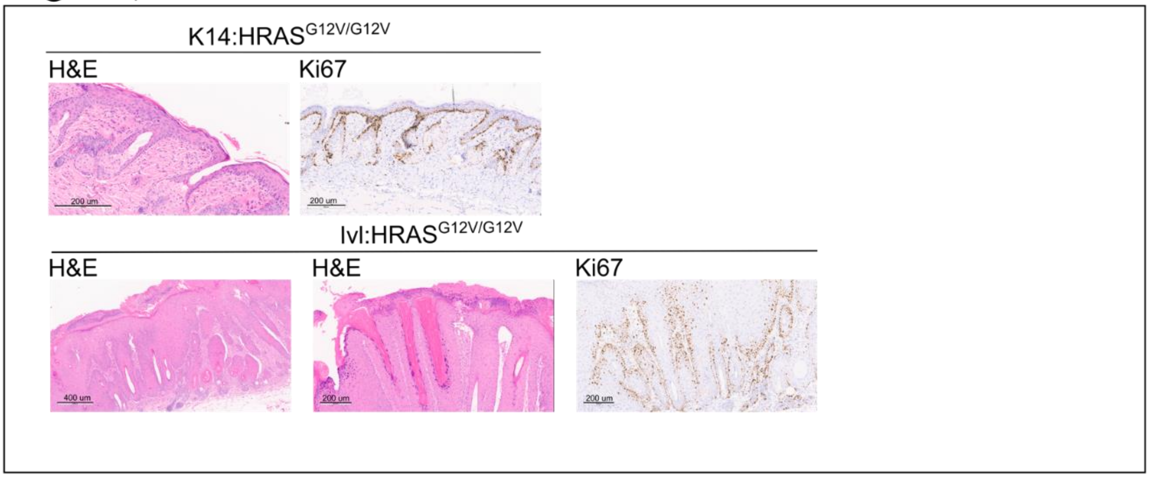


**Figure S1. Histological characterisation of K14:HRAS$^{G12V/G12V}$ skin and Ivl:HRAS$^{G12V/G12V}$ tumour, related to Figure 1**

Represenrative images of H&E and IHC of Ki67 conducted in skin derived from K14:HRAS$^{G12V/G12V}$ (top) and Ivl:HRAS$^{G12V/G12V}$ tumours (bottom) at clinical endpoint.

### Figure S2

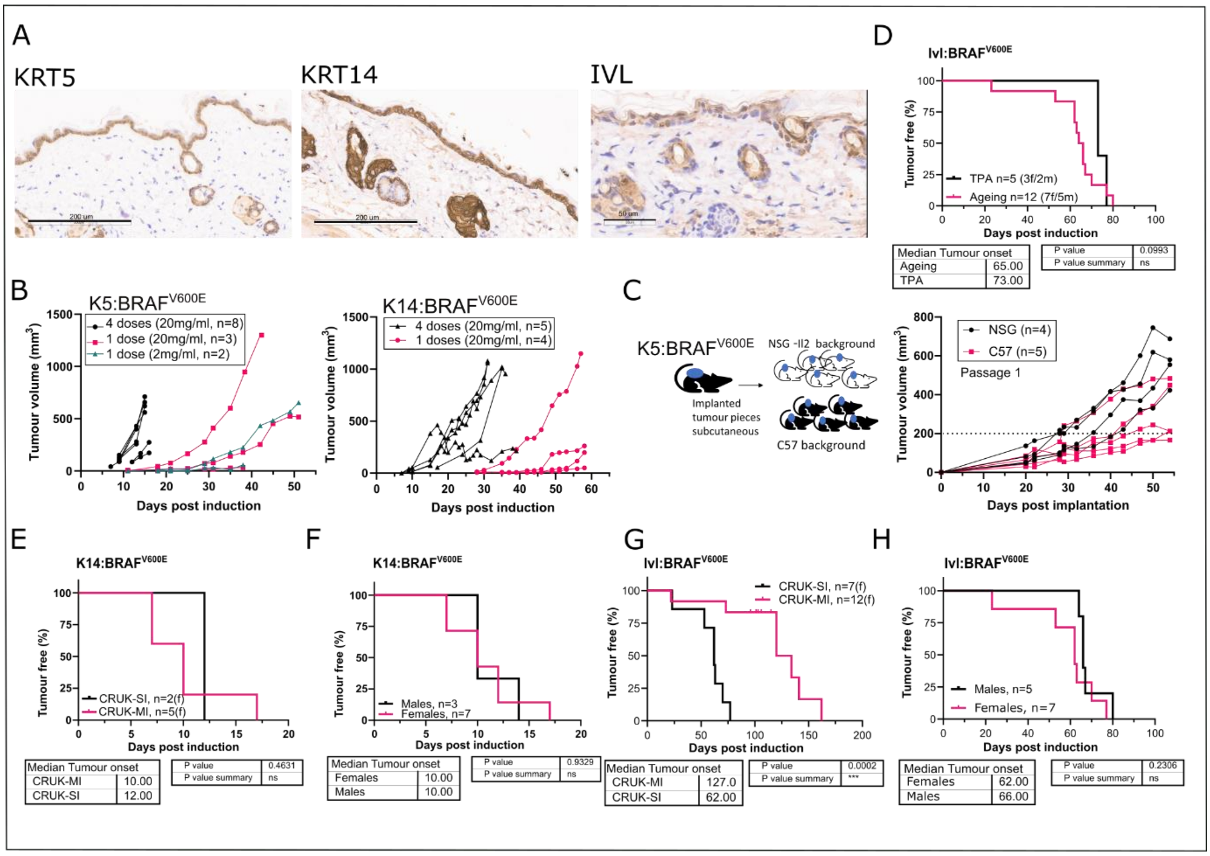


**Figure S2. Tumour-primed and tumour-resistant populations coexist in the basal layer, related to Figure 2**

(A) Representative images of IHC staining of KRT5, KRT14 and IVL in normal wild-type skin.

(B) Tumour burden growth at different induction regimens in K5:BRAF$^{V600E}$ and K14:BRAF$^{V600E}$ models.

(C) Schematic and tumour burden growth of K5:BRAF$^{V600E}$ tumour pieces transplanted into C57/6J and immunosuppressed female mice (NSG-Il2).

(D) Kaplan-Meier tumour-free survival plot for Ivl:BRAF$^{V600E}$ mice, aged until clinical endpoint after treatment with the tumour promoter TPA . P-values were determined using the log-rank (Mantel-Cox) test. Note that the ageing cohort was conducted in CRUK-SI.

(E,F) Kaplan-Meier tumour-free survival plots of K14:BRAF$^{V600E}$ mice, aged until clinical endpoint housed at CRUK Manchester Institute (CRUK-MI) and CRUK Scotland Institute (CRUK-SI) facilities (E, note that CRUK-MI cohort also shown in 2B for comparison), and by sex (F, note that female cohort includes n=5 housed at CRUK-MI, also shown at E, and an additional n=2 housed at CRUK-SI. All males housed at CRUK-SI). P-values were determined using the log-rank (Mantel-Cox) test.

(G,H) Kaplan-Meier tumour-free survival plots of Ivl:BRAF$^{V600E}$ mice, aged until clinical endpoint (G, note that CRUK-MI cohort also shown in 2B for comparison), and by sex (H, all mice housed at CRUK-SI). P-values were determined using the log-rank (Mantel-Cox) test.

### Figure S3

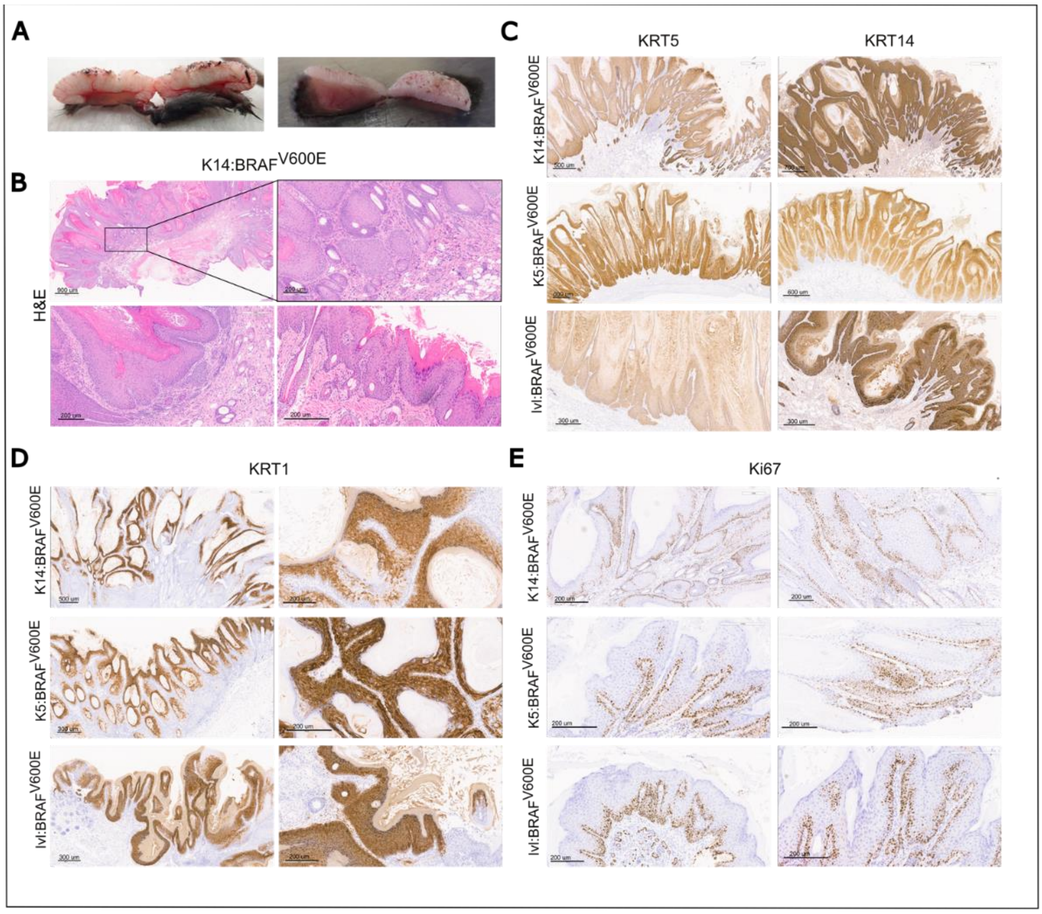


**Figure S3. Histological features of BRAF$^{V600E}$ driven tumours, related to Figure 2**

(A) Representative macroscopic images of the tumours resulting from the tumour-prime population showing demarked borders and vertical columns of keratinocyte proliferation.

(B-E) Representative images of a K14:BRAF$^{V600E}$ tumour showing H&E staining (B), and IHC of skin hierarchy markers KRT5, KRT14 and KRT1 (C and D), and proliferation marker Ki67 (D) at clinical endpoint.

### Figure S4

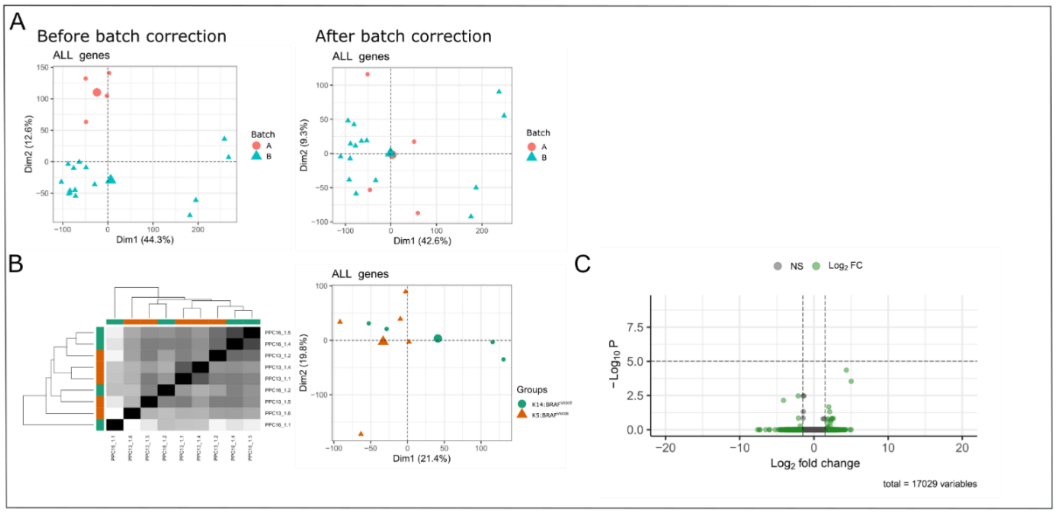


**Figure S4. Transcriptional landscape of tumour-primed K14/K5:BRAF$^{V600E}$ and tumour-resistant Ivl:BRAF$^{V600E}$, related to Figure 4**

(A) Principal component analysis (PCA) of normalised expression values before and after batch correction, showing the correlation between the two different batches sequenced at different times (batch A n=4 and batch B n=15).

(B) Hierarchical clustering heatmap and PCA of normalised expression values showing the correlation between the transcriptomes of the K5:BRAF$^{V600E}$ (n=5) and K14:BRAF$^{V600E}$tumours (n=4) subsets.

(C) Volcano plot showing differentially expressed genes between K5:BRAF$^{V600E}$ and K14:BRAF$^{V600E}$ tumours using Wald test (two-tailed). NS, not significant.

### Figure S5

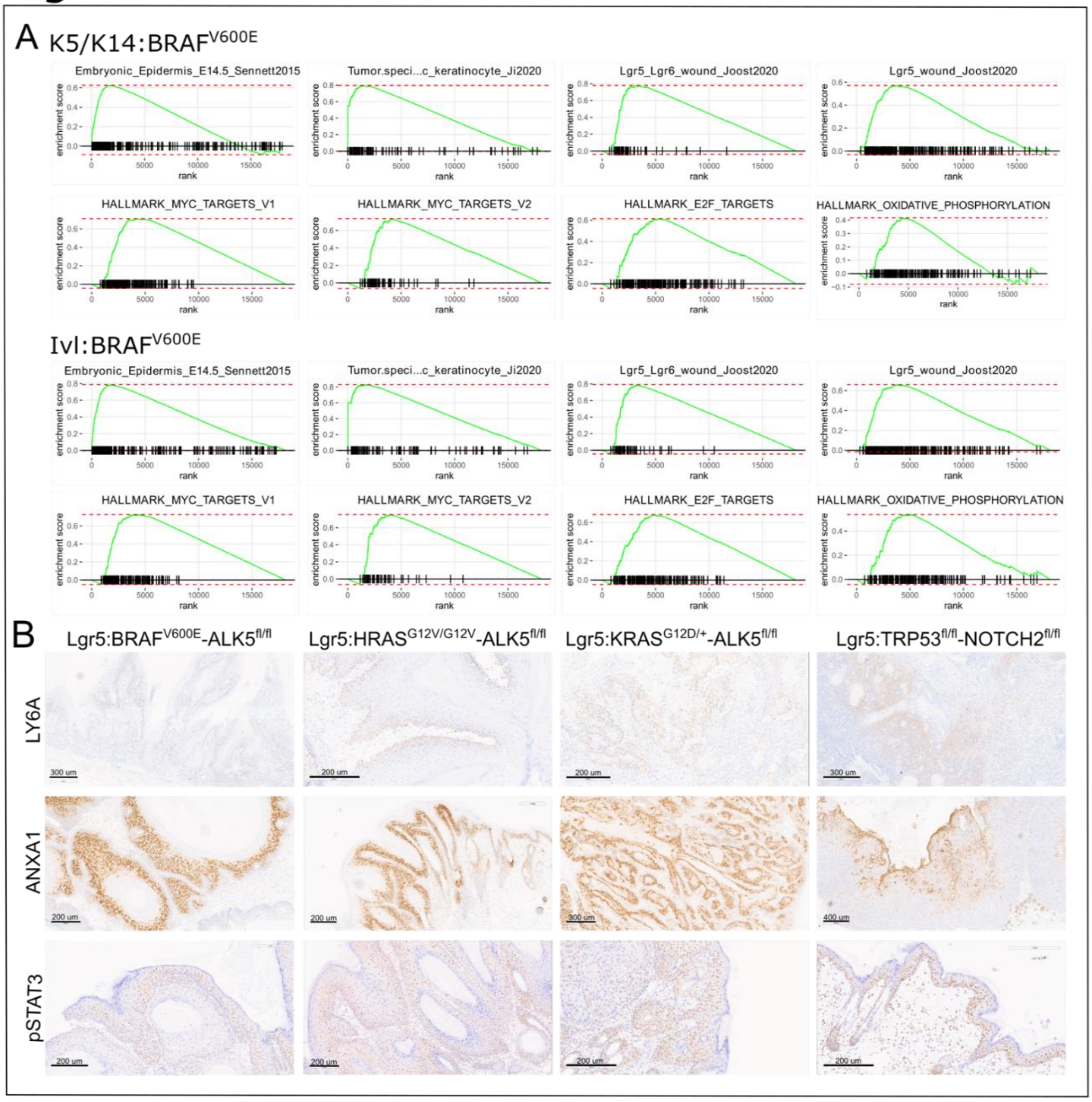


**Figure S5. Shared transcriptional landscape, related to Figure 4**

(A) Selected gene set enriched Hallmarks and pathways in K5/K14:BRAF$^{V600E}$ and Ivl:BRAF$^{V600E}$ tumours at clinical endpoint. Pathways shown are significantly enriched (padj<0.01 based on an adaptive multi-level split Monte-Carlo).

(B) Representative images of IHC validation of selected targets in Lgr5:BRAF$^{V600E}$-ALK5$^{fl/fl}$, Lgr5:HRAS$^{G12V/G12V}$-ALK5$^{fl/fl}$, Lgr5:KRAS$^{G12D/+}$-ALK5$^{fl/fl}$ and Lgr5:TRP53$^{fl/fl}$-NOTCH2$^{fl/fl}$ tumours at clinical endpoint.

Figure S6

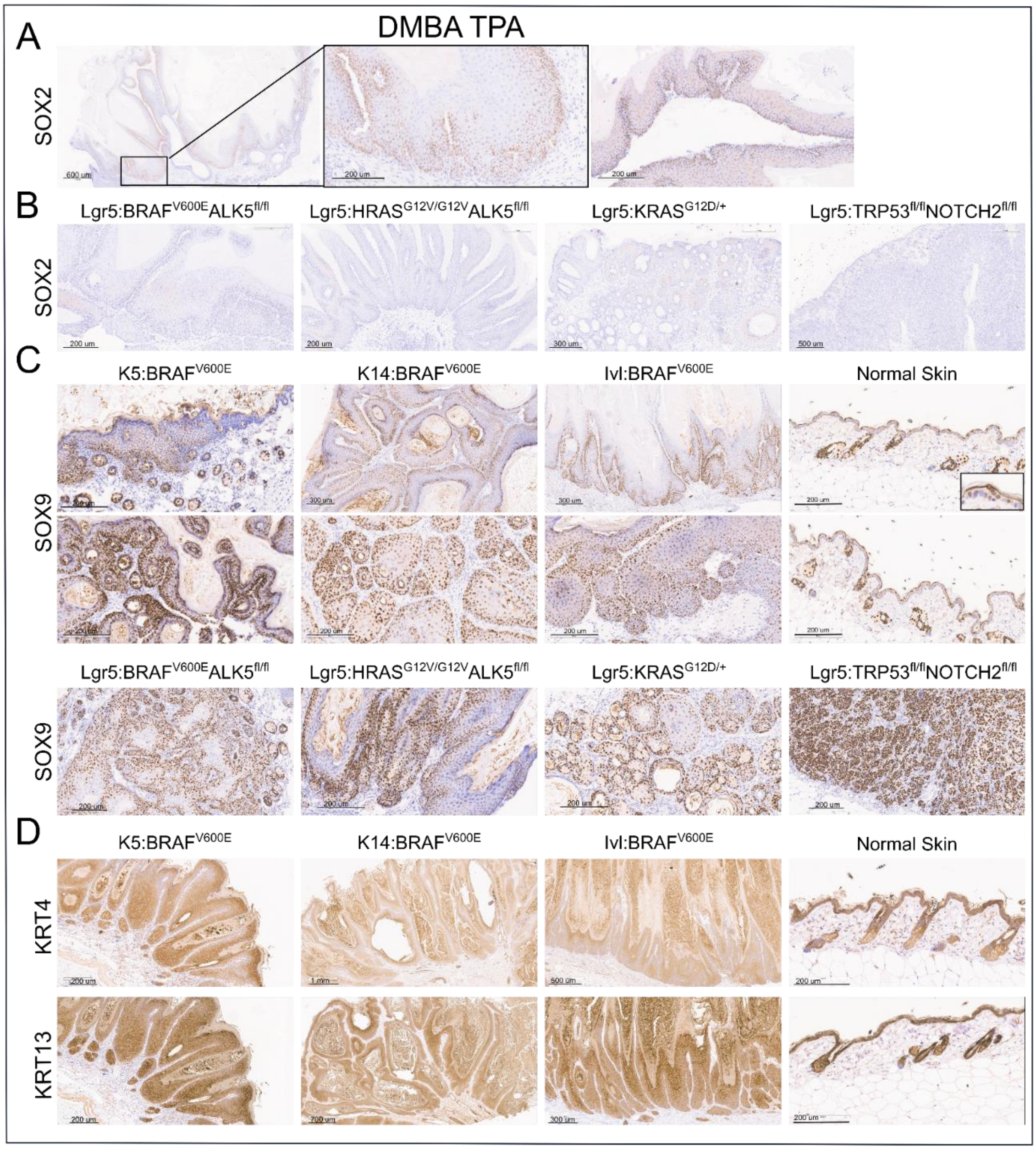


**Figure S6. The tumour-resistant basal population depends on SOX2 for transformation, related to Figure 5**

(A,B) Representative images of IHC of SOX2 in DMBA/TPA-derived tumours (A), and tumours derived from the *Lgr5* population with different driver combinations: Lgr5:BRAF$^{V600E}$-ALK5, Lgr5:HRAS$^{G12V}$-ALK5, Lgr5:KRAS$^{G12D}$-ALK5 and Lgr5:TRP53-NOTCH2 at clinical endpoint (B).

(C) Representative images of IHC of SOX9 in K5:BRAF$^{V600E}$, K14:BRAF$^{V600E}$ and Ivl:BRAF$^{V600E}$ tumours, normal skin and Lgr5:BRAF$^{V600E}$-ALK5$^{fl/fl}$, Lgr5:HRAS$^{G12V/G12V}$-ALK5$^{fl/fl}$, Lgr5:KRAS$^{G12D/+}$-ALK5$^{fl/fl}$ and Lgr5:TRP53$^{fl/fl}$-NOTCH2$^{fl/fl}$ tumours at clinical endpoint.

(D) Representative images of IHC of KRT4 and KRT13 in K5:BRAF$^{V600E}$, K14:BRAF$^{V600E}$ and Ivl:BRAF$^{V600E}$ tumours and in normal skin.

(F) Representative images of IHC of SOX9 in K5:BRAF$^{V600E}$, K14:BRAF$^{V600E}$ and Ivl:BRAF$^{V600E}$ tumours, normal skin and Lgr5:BRAF$^{V600E}$-ALK5$^{fl/fl}$, Lgr5:HRAS$^{G12V/G12V}$-ALK5$^{fl/fl}$, Lgr5:KRAS$^{G12D/+}$-ALK5$^{fl/fl}$ and Lgr5:TRP53$^{fl/fl}$-NOTCH2$^{fl/fl}$ tumours at clinical endpoint.

### Figure S7

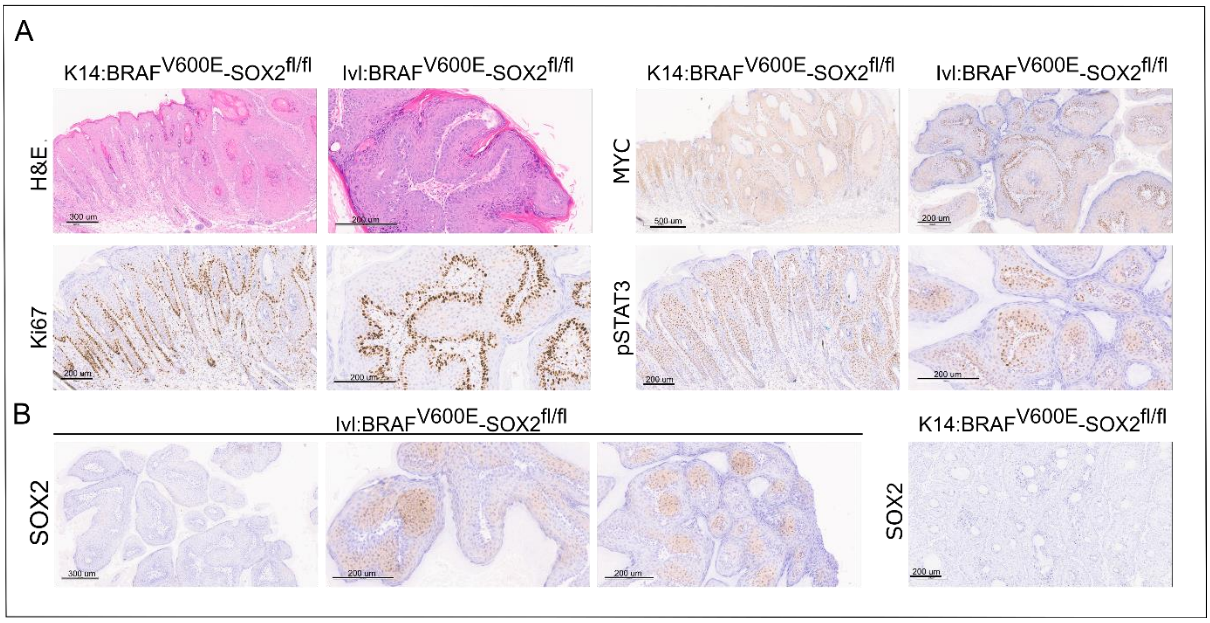


**Figure S7. SOX2 is sufficient to initiate transformation in the tumour-resistant population, related to Figure 6.**

(A) Representative H&E and IHC of Ki67, MYC and pSTAT3 in K14:BRAF$^{V600E}$-SOX2$^{fl/fl}$ and Ivl:BRAF$^{V600E}$-SOX2$^{fl/fl}$ tumour at clinical endpoint.

(B) IHC validation of SOX2 expression in Ivl:BRAF$^{V600E}$-SOX2$^{fl/fl}$ tumours revealing partial expression of SOX2 and no expression in K14:BRAF$^{V600E}$-SOX2$^{fl/fl}$.

### Figure S8

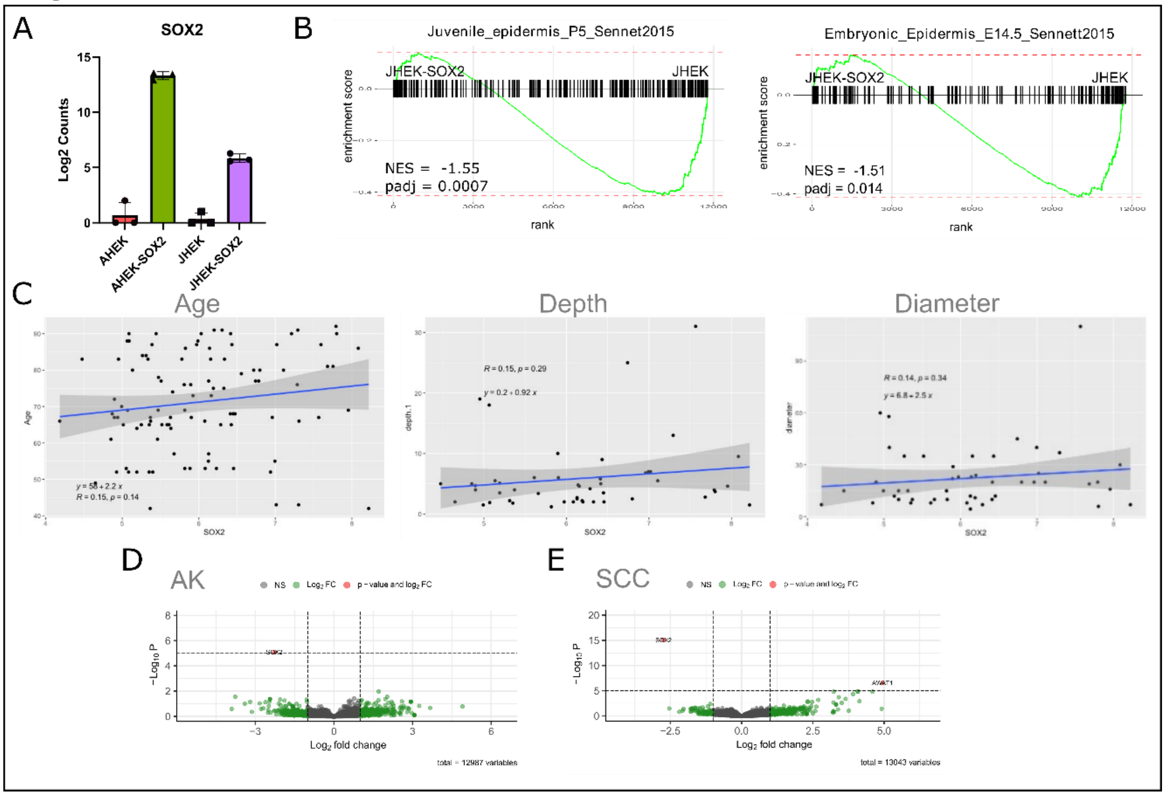


**Figure S8. SOX2 overexpression in juvenile keratinocyte drives a limited transcriptional change, related to Figure 7.**

(A) Normalised RNA expression levels (counts) of SOX2 in AHEK, JHEK, and SOX2-overexpressing AHEK-SOX2 and JHEK-SOX2. N=3, repeated independent experiments per group.

(B) Gene set enrichment analysis of Embryonic epidermis and juvenile transcriptional signatures (Sennett et al., 2015) [23] in JHEK and JHEK-SOX2.

(C) Normalised SOX2 levels from the SCC group [16] correlated with age, depth and diameter.

(D,E) Volcano plot showing differentially expressed genes between SOX2 high and low samples from the AK (D) and SCC (E) groups [16] using the Wald test (two-tailed).
